# Neural correlates of object and action naming practice

**DOI:** 10.1101/590026

**Authors:** Ekaterina Delikishkina, Angelika Lingnau, Gabriele Miceli

## Abstract

Word retrieval deficits are a common problem in patients with stroke-induced brain damage. While complete recovery of language in chronic aphasia is rare, patients’ naming ability can be significantly improved by speech therapy. A growing number of neuroimaging studies have tried to pinpoint the neural changes associated with successful outcome of naming treatment. However, the mechanisms supporting naming practice in the healthy brain have received little attention. Yet, understanding these mechanisms is crucial for teasing them apart from functional reorganization following brain damage. To address this issue, we trained a group of healthy monolingual Italian speakers on naming pictured objects and actions for ten consecutive days and scanned them before and after training. Although activity during object vs. action naming dissociated in several regions (lateral occipitotemporal, parietal and left inferior frontal cortices), training effects for the two word classes were similar and included activation decreases in classical language regions of the left hemisphere (posterior inferior frontal gyrus, anterior insula), potentially due to decreased lexical selection demands. Additionally, MVPA revealed training-related activation changes in the left parietal and temporal cortices associated with the retrieval of knowledge from episodic memory (precuneus, angular gyrus) and facilitated access to phonological word forms (posterior superior temporal sulcus).

## 1. Introduction

Attempted naming can improve performance in aphasic individuals even in the absence of feedback or corrections (Howard, 2000; Nickels, 2002). Although the neural mechanisms underlying this improvement are not clear, it has been recently suggested that they may at least partially overlap with those that support naming facilitation in healthy controls (Heath et al., 2015; Kurland, Liu, & Stokes, 2018). Thus, identifying the neural changes induced by training in healthy participants is necessary to establish a “baseline”, against which the results of anomic patients could be compared.

Studies of incidental naming practice show that even a single instance of naming in the context of a picture naming task can facilitate subsequent processing of a stimulus for days and even weeks (Meister et al., 2005; van Turennout, Bielamowicz, & Martin, 2003; van Turennout, Ellmore, & Martin, 2000). While behaviorally this effect, known as repetition priming, manifests itself as shorter naming latencies, at the neural level it is reflected by decreased activity (or “repetition suppression”) in bilateral occipitotemporal and left prefrontal cortices, associated with facilitated perceptual/conceptual and linguistic processing of a stimulus respectively.

The effects of explicit naming practice were addressed by Basso et al. (2013), who used an intensive training paradigm, more closely resembling speech therapy in patients. Explicit training of object naming (ten repetitions per day over ten consecutive days) was associated with decreased BOLD response in the left inferior frontal cortex and the fusiform gyrus, in line with the studies on repetition suppression, and with increased response in the precuneus and the posterior cingulate cortex. Increased activation of the medial parietal areas, which are not involved in the classic language network, was attributed to retrieval of memories related to practiced items from long-term memory. Similar findings were reported by MacDonald et al. (2015) in healthy older adults, who showed increased activity in the precuneus and decreased activity in the inferior frontal and inferior temporal cortices bilaterally following two object naming sessions (three repetitions/session).

Practice-related activation changes could be modulated by a number of factors (intensity of practice, interval between stimulus repetitions, etc.). Yet, one critical factor, namely, the content of training, has received little attention. Most studies have focused on practiced naming of objects, that are referred to by nouns. Neuropsychological findings (for reviews, see Cappa & Perani, 2003; Mätzig, Druks, Masterson, & Vigliocco, 2009), as well as recent neuroimaging studies with healthy individuals (for reviews, see Crepaldi, Berlingeri, Paulesu, & Luzzatti, 2011; Vigliocco, Vinson, Druks, Barber, & Cappa, 2011) suggest that words belonging to different grammatical classes, such as nouns and verbs, may have at least partially dissociable neural correlates. Thus, it seems reasonable to expect them to be differently affected by practice. A recent study by Kurland et al. (2018) attempted to address this question by including both nouns and verbs in their training protocol, but failed to find a significant interaction between training and word class. It is possible, however, that these null results were due to low intensity of practice (five repetitions a few days prior to the fMRI session, plus five repetitions immediately before the scanning).

In the present fMRI study we investigated differences in the magnitude and localization of training effects for nouns and verbs. Since subjects were instructed to remain silent inside the scanner during picture presentation (in order to avoid jaw movement artifacts) until they saw the next slide with a response cue, we gathered reaction time (RT) data from a separate group of volunteers who participated in an analogous study in which they were asked to produce a word as soon as they saw a picture. Healthy speakers of Italian practiced naming of objects and actions for ten consecutive days and were tested twice, on the days preceding and following the training. The two experimental sessions were identical and included trained items, as well as an equal number of untrained items that served to control for task habituation and priming effects. The use of this paradigm allowed us to investigate: (1) the putative distinctions in the neural representations of objects (nouns) and actions (verbs), (2) the effects of training and their potential interaction with word class, and (3) the reliability of long-term priming effects reported in previous studies (Meister et al., 2005; Meltzer, Postman-Caucheteux, McArdle, & Braun, 2009), separated from the effects of explicit training.

## 2. Materials and Methods

### 2.1. Participants

A total of 35 native Italian speakers took part in this project—12 subjects (3 male, mean age: 23.3 ± 2.5 years, age range: 19-28 years) participated in the behavioral study, and 23 (9 male, mean age: 23.7 ± 3.3 years, age range: 19-32 years) in the fMRI study. Three subjects of the fMRI study were subsequently excluded from data analyses—two because of excessive head motion during scanning (more than 3 mm in one of the directions) and one due to non-compliance with the training protocol. All participants but one were right-handed according to the Edinburgh Handedness Inventory (Oldfield, 1971); the remaining subject was a self-reported right-hander, but scored as ambidextrous on the Inventory. All participants had normal or corrected-to-normal vision and reported no history of neurological or psychiatric disease.

The study was conducted in compliance with the Declaration of Helsinki and was approved by the Human Research Ethics Committee of the University of Trento. All participants signed informed consent forms.

### 2.2. Stimuli

#### 2.2.1. Preliminary naming task

All participants of the behavioral (N = 12) and the fMRI (N = 23) study were required to undergo the preliminary naming task prior to entering the two-session training study, in order to assure that they recognized objects and actions that would be presented in the two experimental sessions and retrieved their corresponding names. Stimuli consisted of line drawings of 80 objects and 80 actions that were presented at a comfortable pace using PowerPoint. Subjects were instructed to produce names of objects using an Italian noun in a singular form (without an article) and to name actions using a verb in the infinitive form. Part of the stimuli were specifically drawn for the present study, while others were selected from various sources, including the Verb and Action Test (VAT; Bastiaanse, Wieling, & Wolthuis, 2016), the Battery for the Analysis of Aphasic Deficits (BADA; Miceli, Laudanna, & Capasso, 2001), as well as the public domain (see examples of drawings in Fig. 1).

**Fig. 1.**
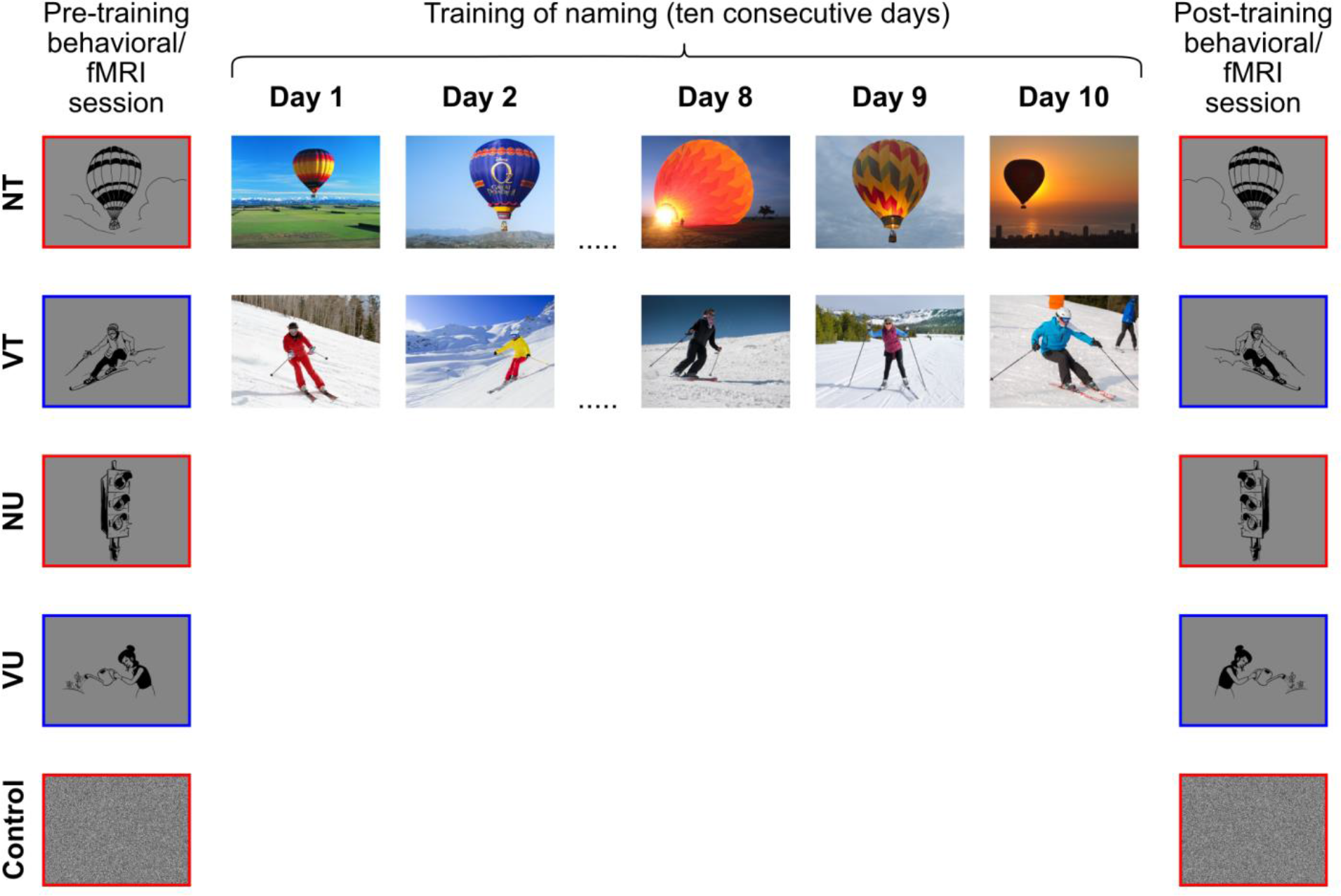
The experimental paradigm. Participants in the behavioral and the fMRI studies named color photographs of 20 objects (NT) and 20 actions (VT) for ten consecutive days. Before and after the training program, they underwent two identical experimental naming sessions that included trained items (NT and VT), as well as an equal number of untrained nouns (NU) and untrained verbs (VU), used as controls for potential stimulus/task habituation effects. Fourier-transformed phase-scrambled images (Control), in response to which subjects were instructed to produce a pseudoword, were also presented in both studies.

#### 2.2.2. Experimental naming task

Of the 160 drawings presented in the preliminary task, 40 object and 40 action pictures were used in two identical experimental naming sessions. Half of the items in each set were included in the training protocol, while the other half were not explicitly trained and served as controls for potential stimulus priming/task habituation effects. The four resulting 20-item subsets—untrained nouns (NU), trained nouns (NT), untrained verbs (VU) and trained verbs (VT)—were matched for variables that reportedly affect word retrieval. Words in the four subsets were balanced for phonemic (*H*(3) = .43, *p* = .934) and syllabic (*H*(3) = .804, *p* = .848) length, as well as for relative lemma frequency (*H*(3) = .006, p = .996), based on a lexical database of written Italian *(Corpus e Lessico di Frequenza dell’Italiano Scritto, CoLFIS*; Bertinetto et al., 2005). Online questionnaires, created on the website SurveyMonkey.com, were delivered to separate groups of Italian native speakers in order to balance target words for familiarity (60 participants; *H*(3) = 2.629, *p* = .452), imageability (38 participants; *H*(3) = 6.549, *p* = .088) and subjective age of acquisition (55 participants; *H*(3) = 3.473, *p* = .324). Another questionnaire was used to balance naming agreement of pictorial stimuli (48 participants; *H*(3) = 2.158, *p* = .54). Additionally, pictures were matched for objective visual complexity (*H*(3) = .245, *p* = .97) using the GIF lossless compression method (Forsythe, Mulhern, & Sawey, 2008). Nouns were selected from a broad range of semantic categories, including animals, professions, clothing, furniture, buildings, vehicles, fruit and vegetables. Verbs were roughly matched for transitivity (VT: 11 transitive, 9 intransitive; VU: 11 transitive, 9 intransitive) and instrumentality (VT: 10 instrumental, 10 non-instrumental; VU: 9 instrumental, 11 non-instrumental). Pictures were normalized with an average brightness of 128 cd/m^2^. Fourier-transformed phase-scrambled images were additionally introduced into the experimental set as low-level controls. In the second experimental session all images were flipped horizontally in order to reduce potential effects of priming in early visual areas.

#### 2.2.3. Training materials

Subjects were asked to practice overt naming of 20 objects in the NT subset and 20 actions in the VT subset using ten booklets with color photographs (one for each day of training). Photos were taken from the public domain and represented various depictions of to-be-trained objects and actions (see Fig. 1 for examples). Each booklet contained a different exemplar of the same concept, in order to tap into abstract structural representations rather than low-level perceptual features. A booklet was divided into two sections—“Objects” and “Actions”. Items within each section were presented in random order.

### 2.3. Procedure

The experimental paradigm is schematically depicted in Fig. 1. Subjects underwent intensive naming training for ten consecutive days (excluding weekends). The training material consisted of 20 objects (NT subset) and 20 actions (VT subset). Training was carried out at home, at a time comfortable for a subject. To make sure that participants complied with the training protocol, they were asked to record their responses with the help of a digital recorder. A daily training session consisted of naming all objects and actions in a given booklet for ten times.

All subjects completed two identical experimental sessions—one before and one after the training, either inside or outside of the MRI scanner, while their RTs were measured. In addition to the 40 trained objects and actions (NT and VT subsets), participants were presented with an equal number of untrained items (NU and VU subsets). Their task was to name a depicted object or action aloud, using a single Italian word (a noun without an article, a verb in the infinitive form). Whenever a scrambled image appeared, subjects were instructed to produce a pseudoword—*/ber.’to:va/* (in session 1) or */sin.’to:ti/* (in session 2).

Stimuli were delivered in blocks. A run consisted of four blocks—NU, NT, VU and VT—presented in random order. Each block included five items belonging to one of the four experimental conditions, as well as two randomly interspersed scrambled images. Word class was cued by a colored frame around an image: a red frame for nouns (NU and NT blocks), a blue frame for verbs (VU and VT blocks). A frame was also placed around scrambled images that appeared within a noun or a verb block. Subjects were instructed to produce the same pseudoword irrespective of frame color. All stimuli (N = 80) were presented within four experimental runs, and subsequently repeated within four additional runs in a different order. Due to technical reasons, for one of the participants of the fMRI study only four out of eight runs were acquired in the first experimental session.

Prior to each experimental session, participants received written instructions and underwent short practice. Stimulus presentation and response collection were controlled with ASF (Schwarzbach, 2011), a toolbox based on Psychtoolbox-3 (Brainard, 1997) for MATLAB (MathWorks, Natick, MA, U.S.A.).

#### 2.3.1. Behavioral naming study (N = 12 participants)

The first experimental session took place on the day after the preliminary naming task. Training started 1 to 3 days after the first experimental session (mean: 1.9 days) and finished on the day preceding the second experimental session. Subjects were allowed to refrain from training during weekends (mean: 2 days). Each trial started with a 2 s black fixation cross followed by a 3 s picture presentation. The inter-trial interval (ITI) was set to 1 s. Blanks of 5 s were introduced between blocks, and at the beginning and the end of each run. Subjects were asked to reply as soon as they saw a picture. Stimuli were presented on an LCD screen with the resolution of 1920 × 1080 pixels and the frame rate of 60 Hz.

#### 2.3.2. fMRI naming study (N = 23 participants)

The preliminary naming task was administered 1 to 4 days prior to the first fMRI session (mean interval: 1.5 days). The training procedure started 1 to 4 days after the first fMRI session (mean: 1.9 days) and finished on the day preceding the second fMRI session. Subjects were allowed to take 1-4 days of rest from training (mean: 2.2 days). Each trial started with a black fixation cross whose presentation lasted between 2 and 5 s. The duration of the initial fixation was chosen from a geometric distribution (*p* = .4; in steps of 1 s). The fixation cross was followed by a picture, presented for 2 s. Subjects were instructed to withhold overt responses while viewing the picture and to respond when a green fixation cross following the picture appeared (3.5 s). The ITI was jittered between 0.5 and 1 s (in steps of 0.25 s). Blanks with a duration of 6 s were introduced between blocks. Each run started and ended with a 12 s blank. In the scanner, stimuli were back-projected onto a screen (frame rate: 60 Hz, screen resolution: 1024 × 768 pixels) via a liquid crystal projector (OC EMP 7900, Epson, Nagano, Japan). Participants viewed the screen binocularly through a mirror mounted on the head coil.

### 2.4. Data acquisition

#### 2.4.1. Behavioral data acquisition

Vocal responses of the participants of the behavioral study were collected using the Samson Q4 microphone with a low-noise microphone cable (Thomann, Burgebrach, Germany). RTs were measured automatically using the voice key function supplied with ASF. Recordings were digitized at a sampling rate of 44.1 kHz.

#### 2.4.2. MR data acquisition

Neuroimaging data were collected at the Functional Neuroimaging Laboratories (LNiF) of the Center for Mind/Brain Sciences (CIMeC) at the University of Trento, Italy, using a 4T Bruker MedSpec MR scanner with an 8-channel birdcage head coil. Functional images were acquired using a T2*-weighted gradient echo-planar imaging (EPI) sequence with fat suppression. Scanning was performed continuously during a functional run with the following parameters: repetition time (TR) = 2.2 s, echo time (TE) = 33 ms, flip angle (FA) = 75°, field of view (FOV) = 192 × 192 mm, matrix size = 64 × 64, voxel resolution = 3 × 3 × 3 mm. We acquired 31 slices in ascending-interleaved odd-even order, with a thickness of 3 mm and a 15% gap (0.45 mm). Slices were aligned to the AC-PC plane. An imaging volume was positioned to cover the entire temporal lobe; as a result, a small portion of the superior parietal cortex was not captured in most subjects. The number of volumes in a functional run varied (range: 130-142) as a result of temporal jittering introduced into trials. Before each run we performed an additional scan measuring the point-spread function (PSF) of the acquired sequence, in order to correct the distortion in geometry and intensity expected with high-field imaging (Zaitsev, Hennig, & Speck, 2004; Zeng & Constable, 2002). A T1-weighted structural scan at the beginning of each scanning session served as reference for coregistration of functional data. Structural images were acquired using a magnetization-prepared rapid-acquisition gradient echo (MPRAGE) sequence (TR = 2.7 s, TE = 4.18 ms, FA = 7°, FOV = 256 × 224 mm, 176 slices, inversion time (TI) = 1020 ms), with generalized autocalibrating partially parallel acquisition (GRAPPA) with an acceleration factor of 2.

### 2.5. Data analysis

#### 2.5.1. Behavioral data analysis

Voice onset intensity threshold was calibrated for each subject based on the visual inspection of the wave plots of vocal responses with displayed RTs at a given threshold produced by the ASF software for each trial. RTs deviating from a subject’s mean by more than two standard deviations were considered outliers and removed from the analysis (5.2% of the data removed, including 3.4% of object trials, 9.9% of action trials and 1.4% of control trials). After calculating individual descriptive statistics in MATLAB R2015b, the data were submitted to inferential analysis with repeated-measures ANOVAs and paired-samples *t-*tests in SPSS 24.

#### 2.5.2. fMRI data analysis

Neuroimaging data were preprocessed and analyzed using BrainVoyager QX 2.8.4 (Brain Innovation, Maastricht, the Netherlands) in combination with the NeuroElf toolbox (v. 1.1) and in-house software written in MATLAB. The first three volumes of a functional run were discarded to avoid T1 saturation. For each run, we performed slice timing correction (cubic spline interpolation), followed by 3D motion correction (trilinear interpolation for estimation and sinc interpolation for resampling, all functional volumes acquired in a session realigned to the first volume of the first run) and temporal high-pass filtering with linear trend removal (cut-off frequency of 3 cycles per run). For univariate analyses, functional data were spatially smoothed with a Gaussian filter of 6 mm full width at half maximum (FWHM), in order to reduce noise and minimize intersubject anatomical differences. Functional and structural data were aligned in several steps, using the rigid-body transformation with 6 parameters (3 translations, 3 rotations): the first volume of the first functional run in a session was coregistered to an anatomical image for the corresponding session; then, anatomical scans obtained in the two sessions with a participant were aligned to each other; finally, functional data from both sessions were coregistered to one of the anatomical images using the transformation parameters obtained during the intersession anatomical alignment. For group analysis, structural and functional data were standardized to the Talairach stereotactic space (Talairach & Tournoux, 1988), using sinc interpolation.

Statistical analyses were performed with a general linear model (GLM), as implemented in BrainVoyager. A trial was modeled as an epoch lasting from the onset to the offset of a picture (2 s). Regressors included predictors of the 10 experimental conditions (2 sessions × 5 conditions: S1_NU, S1_NT, S1_VU, S1_VT, S1_Control, S2_NU, S2_NT, S2_VU, S2_VT, S2_Control). Additionally, 6 parameters resulting from head motion correction were included in the model as regressors of no-interest. Each predictor was convolved with a dual-gamma hemodynamic response function (HRF; Friston et al., 1998). The resulting reference time courses were used to fit the signal time courses in each voxel.

Analyses were performed on the cortical surface, with the help of cortex-based alignment (CBA) as implemented in BrainVoyager. This procedure enables better alignment of structural and functional data across subjects by taking into account individual variability in gyral and sulcal folding patterns. To this end, we segmented the white/gray matter boundary on individual Talairach-transformed T1-weighted anatomical scans and reconstructed 3D hemispheric meshes for each participant. Then we inflated each mesh to a sphere with cortical curvature maps projected onto it (with four coarse-to-fine levels of smoothing) and aligned it to a standard spherical surface using a coarse-to-fine moving target approach (Fischl, Sereno, Tootell, & Dale, 1999; Goebel, Esposito, & Formisano, 2006). The resulting transformation matrices were used to create group-averaged surface meshes for the left and the right hemisphere. Statistical analyses were performed separately for each hemisphere. Thresholded statistical maps obtained for a group were projected onto the group-averaged hemispheric meshes for visualization and described using the CBA-transformed macroanatomical surface atlases supplied with BrainVoyager.

##### 2.5.2.1. Univariate analysis

For the univariate analysis, we created mesh time courses for each run by sampling the spatially smoothed functional data from −1 to 2 mm from the reconstructed white/gray matter boundary. For the first level analysis (prior to CBA), we ran individual fixed-effects (FFX) GLMs on the subject data collapsed across runs and obtained *t-*statistics for main effects of the experimental conditions. These *t-*maps were subsequently aligned to group-averaged meshes using the aforementioned transformation matrices. At the group level, individual CBA-transformed *t*-maps were submitted to nonparametric permutation testing (Nichols & Holmes, 2002), in combination with Threshold-Free Cluster Enhancement (TFCE; Smith & Nichols, 2009) as implemented in the CoSMoMVPA toolbox (Oosterhof, Connolly, & Haxby, 2016). A total of 1000 Monte Carlo simulations and a corrected cluster threshold *α* = .05 (two-tailed; *z* > 1.96) were used.

##### 2.5.2.2. MVPA

In addition to the standard GLM, we carried out multivariate pattern analysis (MVPA; Haxby et al., 2001). Rather than contrasting the amplitude of the BOLD response in individual voxels/surface vertices, MVPA is based on comparing spatial patterns of activation in response to different experimental conditions (for review, see Haxby, 2012). Specifically, we employed a whole-brain searchlight analysis, a recently developed MVPA technique for identifying locally informative areas of the brain (Etzel, Zacks, & Braver, 2013; Kriegeskorte, Goebel, & Bandettini, 2006), which may outperform mass-univariate analyses due to its greater sensitivity to distributed coding of information (Davis et al., 2014; Jimura & Poldrack, 2012). We performed a searchlight analysis on the brain surface (Oosterhof, Wiestler, Downing, & Diedrichsen, 2011), using a linear discriminant analysis (LDA) classifier as implemented in CoSMoMVPA. We aimed to examine in which areas the classifier could reliably (i.e., significantly above chance) distinguish: (1) nouns vs. verbs (using data from the pre-training fMRI session), and (2) trained vs. untrained items (based on data from the post-training fMRI session). To this end, we ran single-study GLMs separately for each run of each subject, using unsmoothed mesh time courses. At the single-subject level (prior to CBA), maps containing *t*-statistics for the main effects of experimental conditions for each run in a session were stacked together and submitted to the searchlight analysis across the entire cortex using spheres with an 8-mm radius. Classification accuracies were obtained using a leave-one-out cross-validation method with an 8-fold partitioning scheme. The dataset was split into 8 chunks (each corresponding to one experimental run), and the classifier was trained on the data from 7 chunks and tested on the remaining one. The procedure was repeated for 8 iterations, using all possible train/test partitions, and the average decoding accuracies across these iterations were calculated. Decoding accuracies obtained for a given searchlight were assigned to its central vertex. Individual surface maps containing average decoding accuracies were aligned to the group-averaged mesh using the transformation matrices obtained during CBA. At the group level, a two-tailed one-sample *t*-test across individual maps identified vertices where classification was significantly above chance (50%, since our classifiers were binary). The resulting maps were corrected using TFCE with 1000 Monte Carlo simulations (corrected cluster threshold *α* = .05, two-tailed; *z* > 1.96).

## 3. Results

### 3.1. Behavioral results

Average response latencies of the 12 participants in the behavioral experiment are presented in Fig. 2. A three-way repeated-measures ANOVA with session (first, second), word class (noun, verb) and training (trained, untrained) as within-subject factors was carried out. Significant main effects of word class (faster RTs to nouns than verbs; *F*(1, 11) = 68.89, *p* < .001), training (faster RTs to trained than untrained items; *F*(1, 11) = 41.22, *p* < .001) and session (faster RTs in session 2 than in session 1; *F*(1, 11) = 6.32, *p* = .029) were found. The session-by-training interaction was also significant (*F*(1, 11) = 10.57, *p* = .008)—an expected outcome, considering that prior to training untrained and to-be-trained items were indistinguishable. The lack of significant training-by-word-class (*p* = .199, ns) and session-by-word-class (*p* = .285, ns) interactions suggests the magnitude of training and session effects was similar for words belonging to both classes.

**Fig. 2.**
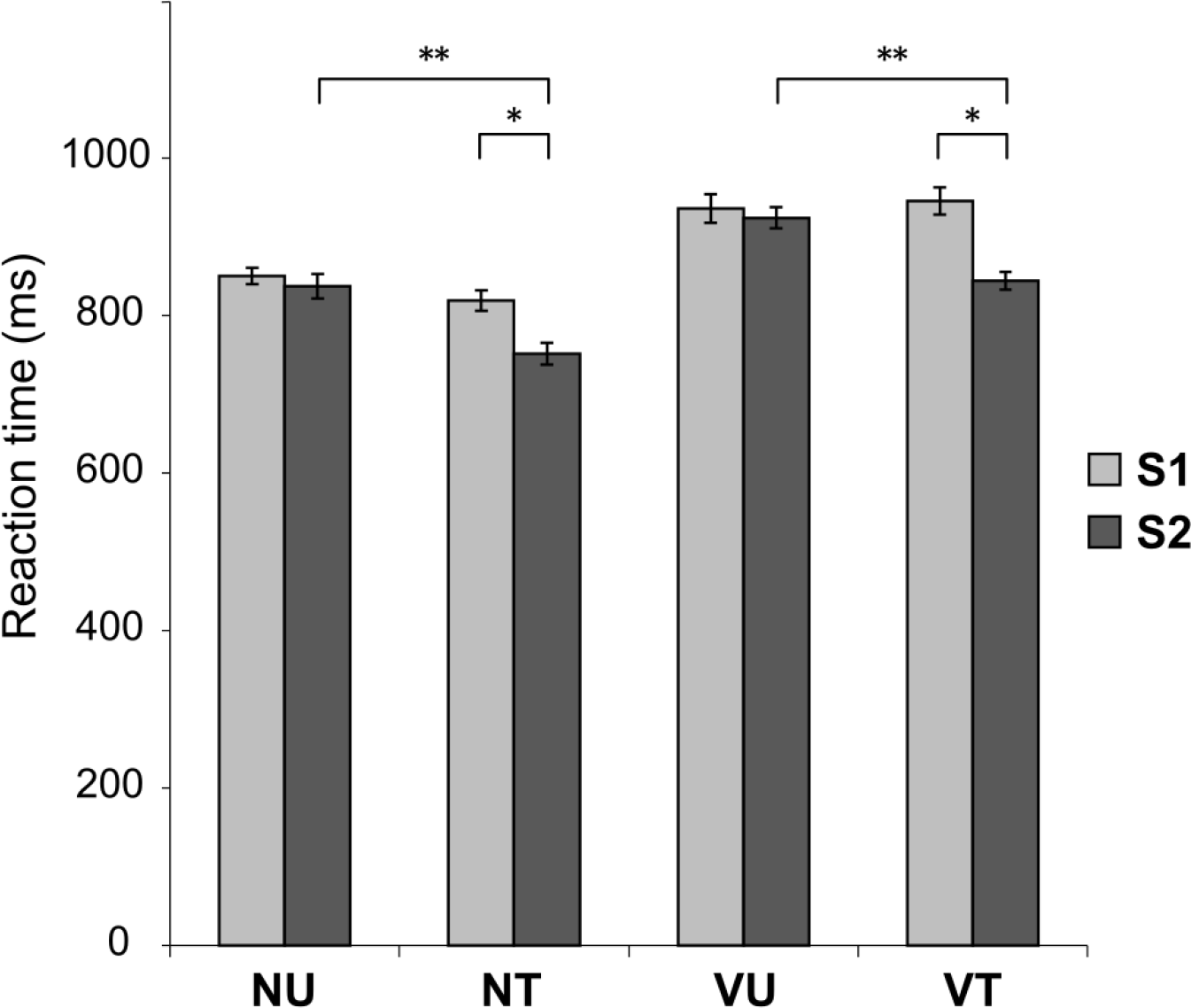
Behavioral results. Average reaction times for each stimulus category in session 1 (S1; light gray bars) and session 2 (S2; dark gray bars). NU = untrained nouns, NT = trained nouns, VU = untrained verbs, VT = trained verbs. Error bars reflect standard errors of the mean after removing between-subject variability (Cousineau, 2005); * denotes significant effects at *p*_FDR_ < .05; ** denotes significant effects at *p*_FDR_ < .005.

To examine whether shorter RTs in the second session were driven mainly by session or by training effects, we conducted six paired-samples *t*-tests comparing trained and untrained items belonging to the same word class within and across sessions. In addition, to rule out significant differences between subsets prior to training, we carried out a *t*-test comparing items from the trained and the untrained subset in session 1 for each word class. The resulting *p*-values were corrected using the false discovery rate (FDR) method for the overall number of comparisons (N = 8; Benjamini & Yekutieli, 2001). The *t*-tests designed to compare the two subsets of nouns and verbs prior to training failed to distinguish RTs to to-be-trained and not-to-be-trained items (nouns: S1_NT vs. S1_NU: *p* = .154, ns; verbs: S1_VT vs. S1_VU: *p* = .611, ns). Significant effects of training, for both nouns and verbs, were observed when comparing responses to trained items before and after training (S2_NT vs. S1_NT: *p*_*FDR*_ = .026; S2_VT vs. S1_VT: *p*_*FDR*_ = .01), and responses to trained and untrained items after training (S2_NT vs. S2_NU: *p*_*FDR*_ = .003, S2_VT vs. S2_VU: *p*_*FDR*_ = .004). However, no significant session effects were found for untrained nouns (S2_NU vs. S1_NU: *p*_*FDR*_ = .694, ns) and verbs (S2_VU vs. S1_VU: *p*_*FDR*_= .645, ns), suggesting that the significant main effect of session had been actually driven by the training effect.

### 3.2. fMRI results

#### 3.2.1. Word class effects

##### 3.2.1.1. Univariate results

Naming of pictured objects and actions activated similar brain networks (Fig. 3 A), as shown by contrasting nouns/verbs with phase-scrambled controls in the first fMRI session (S1_NU + S1_NT > S1_Control; S1_VU + S1_VT > S1_Control). Activations were observed bilaterally in ventral and lateral occipitotemporal areas, including inferior and middle occipital gyri, fusiform and parahippocampal gyri, i.e., in the ventral visual processing stream involved in object and shape recognition (Goodale & Milner, 1992, 2018; Ungerleider & Mishkin, 1982). They extended bilaterally into the posterior portion of the superior parietal lobule (SPL) and anterior insular cortices. Additionally, picture naming recruited most of the left inferior frontal gyrus (IFG), as well as the ventral precuneus and pre-supplementary motor area (pre-SMA). Detailed information about the areas activated in this and other univariate contrasts is provided in Supplementary Table 1.

**Fig. 3.**
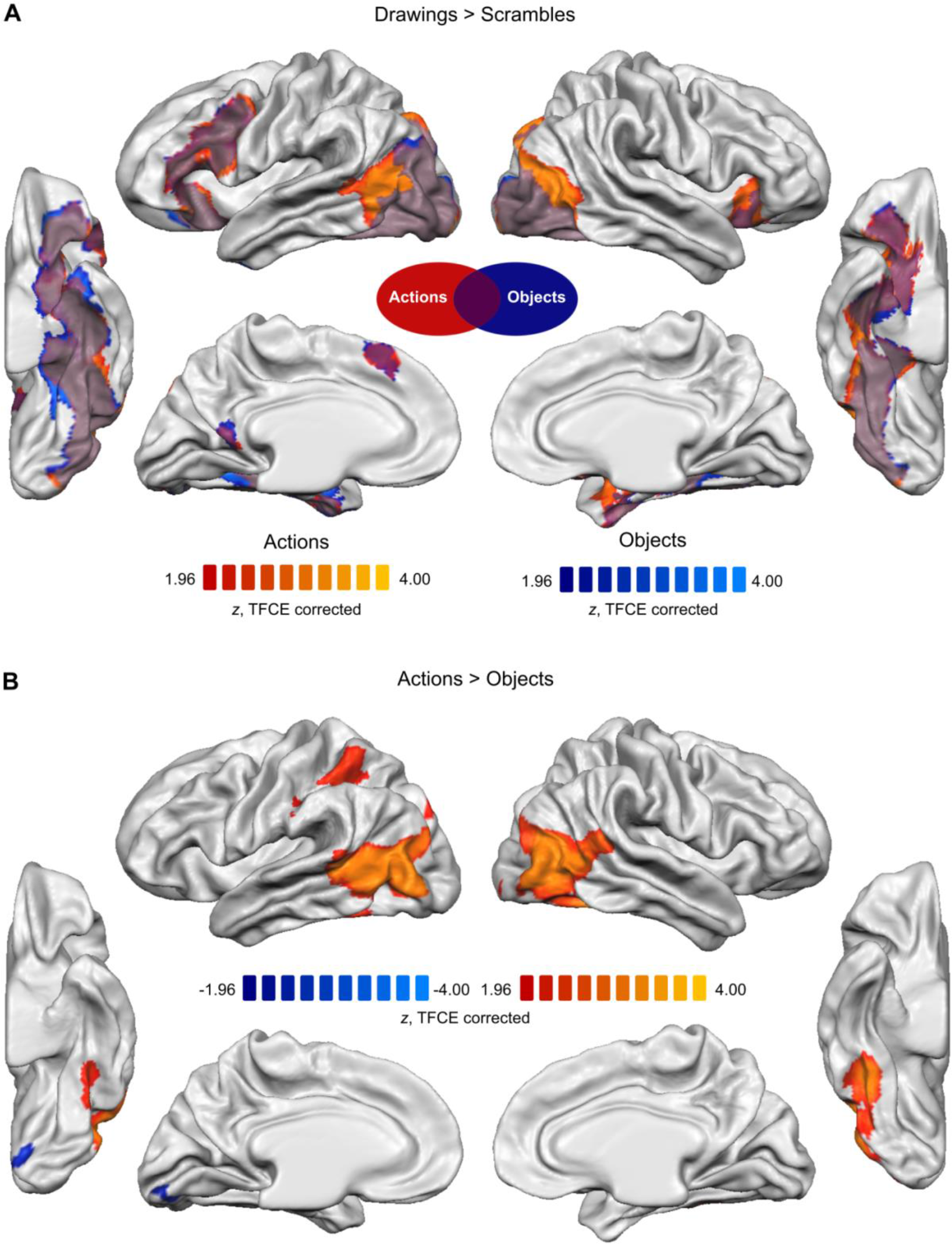
Word class effects, revealed by the univariate RFX GLM contrasts of stimuli presented in the pre-training fMRI session. **A.** Object (blue) and action (red/orange) naming networks, identified by contrasting all nouns/verbs with meaningless scrambled images (negative tail not shown). The logical conjunction of the two maps (i.e., areas recruited both during object and action naming) is indicated in purple. **B.** Areas showing increased BOLD response to verbs as compared to nouns (red/orange) and to nouns as compared to verbs (blue). The statistical group maps (N = 20) for each hemisphere were corrected for multiple comparisons using TFCE (α = .05, two-tailed) and projected onto the group-averaged surface meshes for visualization.

Inspection of Fig. 3 A suggests that bilateral activations associated with verb production were more extensive than those associated with noun production in posterior middle temporal gyri (pMTG) and SPL. The direct contrast of responses to nouns and verbs in session 1 (S1_VU + S1_VT > S1_NU + S1_NT; Fig. 3 B) revealed that verb naming engaged to a significantly greater extent the lateral occipitotemporal cortices (LOTC), as well as regions in the left SPL and the intraparietal sulcus. The opposite contrast (S1_NU + S1_NT > S1_VU + S1_VT) detected a stronger BOLD response for nouns than verbs only in a small cluster in the posterior portion of the medial fusiform/extrastriate cortex.

##### 3.2.1.2. MVPA results

We trained the classifier on data from the pre-training fMRI session to identify the areas in which it would reliably distinguish nouns and verbs. The whole-brain searchlight analysis (Fig. 4) showed that patterns of *t*-scores for nouns and verbs were decodable in a number of bilateral areas, including the LOTC and the SPL, confirming findings of the univariate analysis. Nouns and verbs were also distinguishable on the basis of their patterns of activation in ventral occipitotemporal cortices, early visual areas and the precuneus. Finally, we were able to decode nouns and verbs in the left IFG, extending dorsally into the middle frontal gyrus and caudally into the premotor cortex. Detailed information about the areas identified by the multivariate analyses is provided in Supplementary Table 2.

**Fig. 4.**
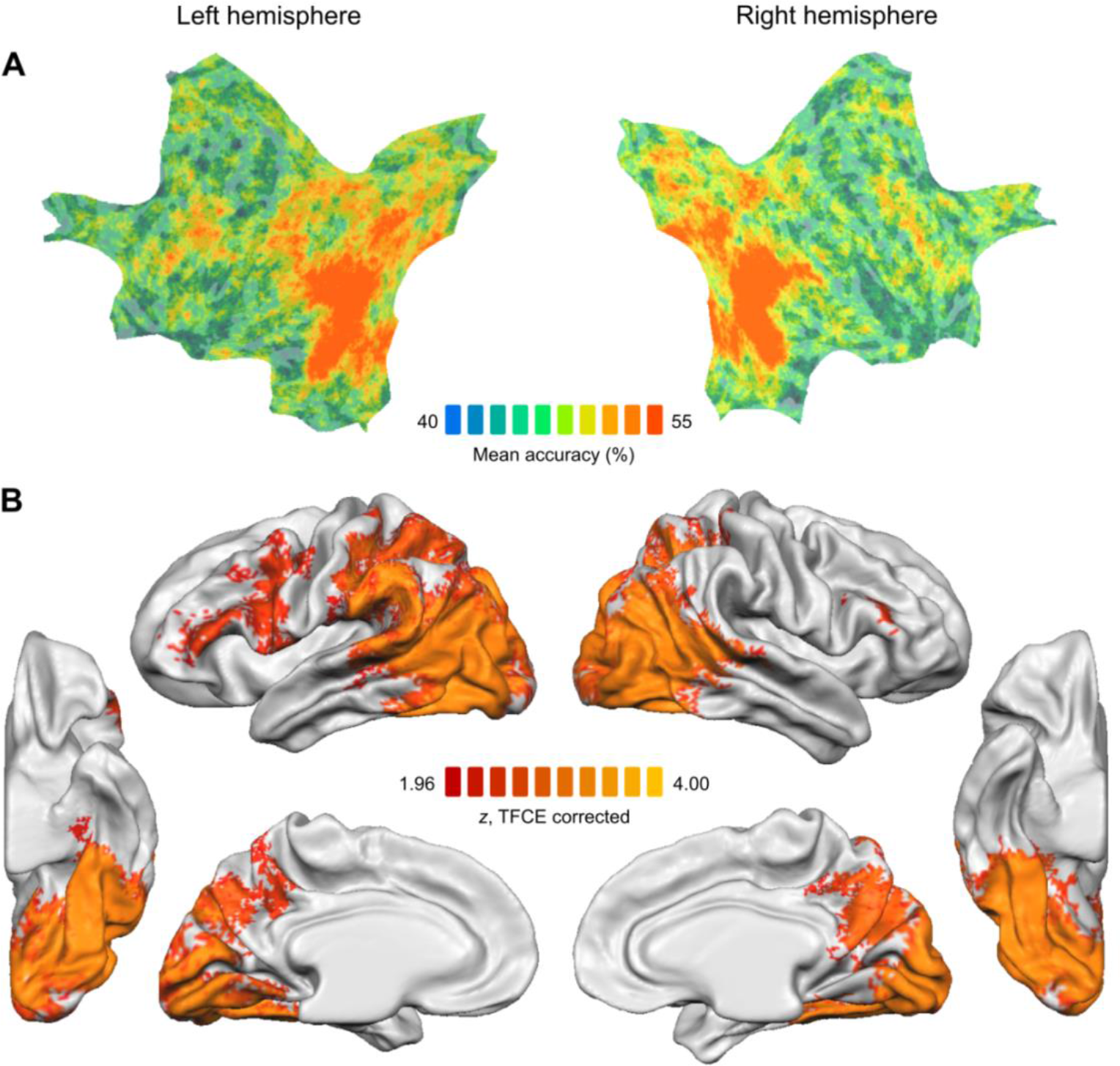
Decoding of word classes (nouns vs. verbs), based on the pre-training fMRI session. **A.** Mean accuracy maps of the searchlight MVPA. Individual accuracy maps (N = 20) were averaged and projected onto a flattened group-averaged hemispheric surfaces. Chance accuracy is 50%. **B.** Statistical group maps, corrected using TFCE at α = .05 (two-tailed).

#### 3.2.2. Training effects

##### 3.2.2.1. Univariate results

To identify the neural correlates of intensive naming practice, we compared responses to trained and untrained items. No significant differences were found when contrasting these items in the first, pre-training fMRI session (S1_NT > S1_NU; S1_VT > S1_VU), attesting to the fact that the two subsets of nouns and the two subsets of verbs had been matched for the relevant variables. In the second, post-training fMRI session, contrasting the same items (S2_NT > S2_NU; S2_VT > S2_VU) revealed significant BOLD amplitude changes in several brain regions (Fig. 5 A). When compared to untrained items in the same post-training session, both trained nouns (Fig. 5 A, left panel) and trained verbs (Fig. 5 A, right panel) yielded a significantly reduced BOLD response in anterior regions of the left hemisphere, including the posterior IFG (pars opercularis and pars triangularis) and the adjacent frontal operculum/anterior insula. Even though deactivations seemed more extensive for verbs than for nouns, the compound contrast (S2_VT > S2_VU) > (S2_NT > S2_NU) failed to reach significance.

**Fig. 5.**
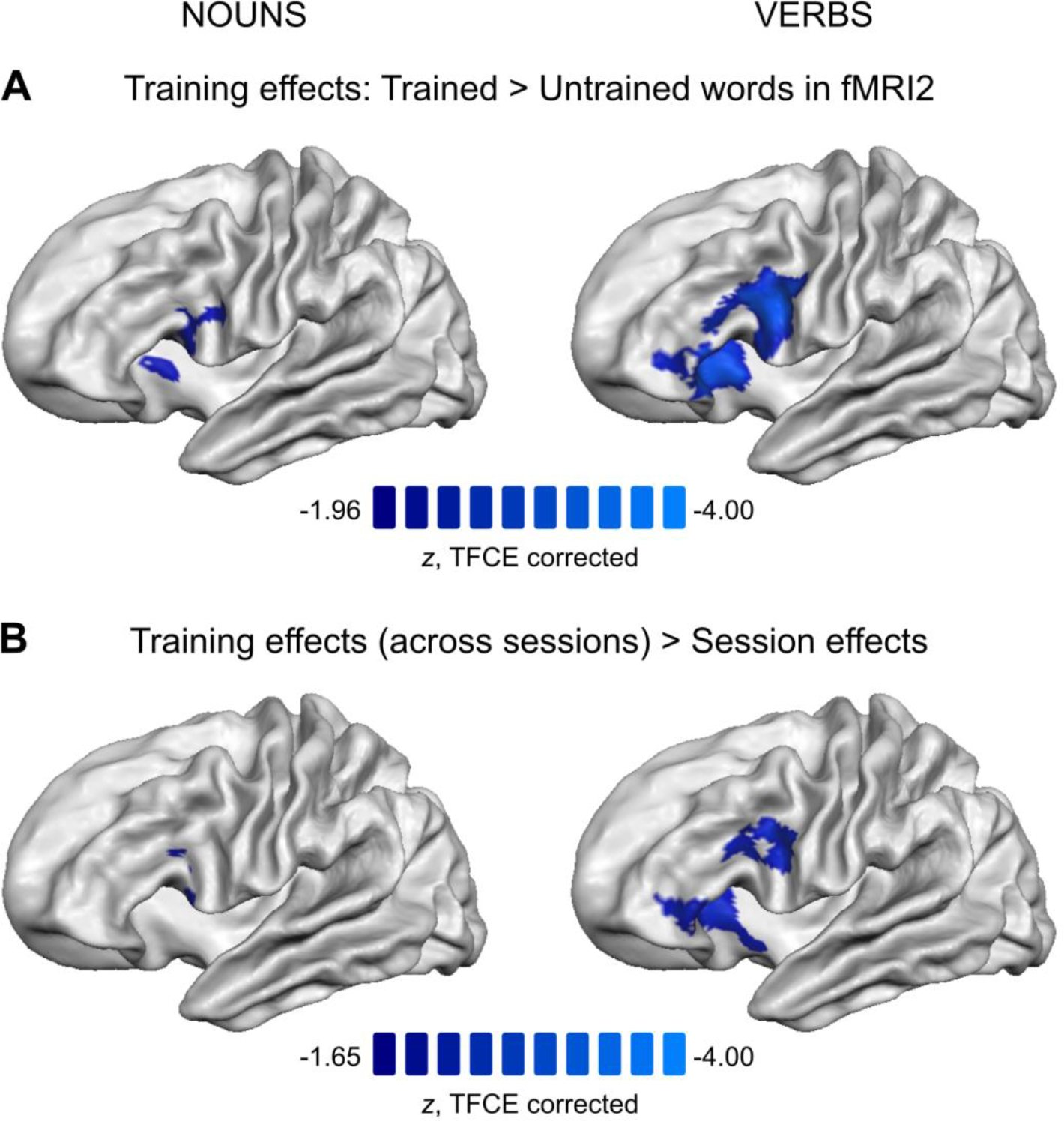
Training effects, revealed by the univariate RFX GLM analysis (N = 20), projected onto the group-averaged mesh of the left hemisphere. **A.** Areas showing significant deactivation following training of nouns (left panel) and verbs (right panel), as evidenced by contrasting words from trained and untrained subsets in the post-training fMRI session (corrected using TFCE at α = .05, two-tailed). **B.** Comparison of training and session effects across the two fMRI sessions (corrected using TFCE at α = .05, one-tailed), separately for nouns (left panel) and for verbs (right panel).

Additionally, we examined the effects of incidental word repetition (i.e., session effects), in order to subsequently compare them with the effects of explicit naming practice (i.e., training effects). By contrasting untrained items in the post- and pre-training sessions (Supplementary Fig. 1), we found that the mere exposure to the same stimuli and the same task (in the same scanner environment) twice over the course of two weeks yielded significant repetition suppression in early visual areas and the fusiform, both for nouns (S2_NU > S1_NU; Supplementary Fig. 1 A) and for verbs (S2_VU > S1_VU; Supplementary Fig. 1 B), in line with reports on priming of low-level features and amodal structural representations respectively (for reviews, see Henson, 2003; Schacter & Buckner, 1998). Decreased activity in the SPL (bilateral for nouns, right-lateralized for verbs) was not expected, but could be explained by facilitated visuospatial processing of familiar stimuli (Beauchamp, Petit, Ellmore, Ingeholm, & Haxby, 2001; Corbetta & Shulman, 1998; Nobre et al., 1997). Additionally, a significantly reduced BOLD response for nouns was observed in the left posterior superior frontal gyrus (on the lateral surface, adjacent to the precentral gyrus). The compound contrast (S2_VU > S1_VU) > (S2_NU > S1_NU) failed to reveal significantly different session effects for the two word classes.

Finally, we directly compared the across-session training and session effects (Fig. 5 B). Since we hypothesized that the BOLD amplitude would decrease more for trained items in the areas associated with explicit practice, we used a one-tailed *t*-test (*z* < −1.65). Indeed, verb training was associated with significantly stronger decrease of the BOLD signal in the left posterior IFG, including a cluster encompassing most of the pars opercularis and a portion of the pars triangularis, as well as clusters in the pars orbitalis and the anterior insula ((S2_VT > S1_VT) > (S2_VU > S1_VU); Fig. 5 B, right panel). An analogous contrast for nouns ((S2_NT > S1_NT) > (S2_NU > S1_NU); Fig. 5 B, left panel) revealed greater training-related reductions of the BOLD response in the left pars triangularis and the adjacent frontal operculum.

##### 3.2.2.2. MVPA results

To further localize areas sensitive to training, we performed searchlight pattern classification analysis on data from the post-training fMRI session. As a first step, the two classifiers learned to distinguish between trained and untrained items, separately for nouns and verbs. Average decoding accuracies did not go beyond chance in any brain region, possibly due to insufficient statistical power. In order to increase the power, the data were collapsed across word classes and a binary classifier was trained to distinguish between trained and untrained items, irrespective of word class. Results (Fig. 6) show that several areas in the left hemisphere (posterior superior temporal sulcus, angular gyrus, precuneus) were sensitive to training. Decoding was also significantly above chance in the left anterior insula (replicating the univariate results) and in two small clusters close to the right calcarine sulcus.

**Fig. 6.**
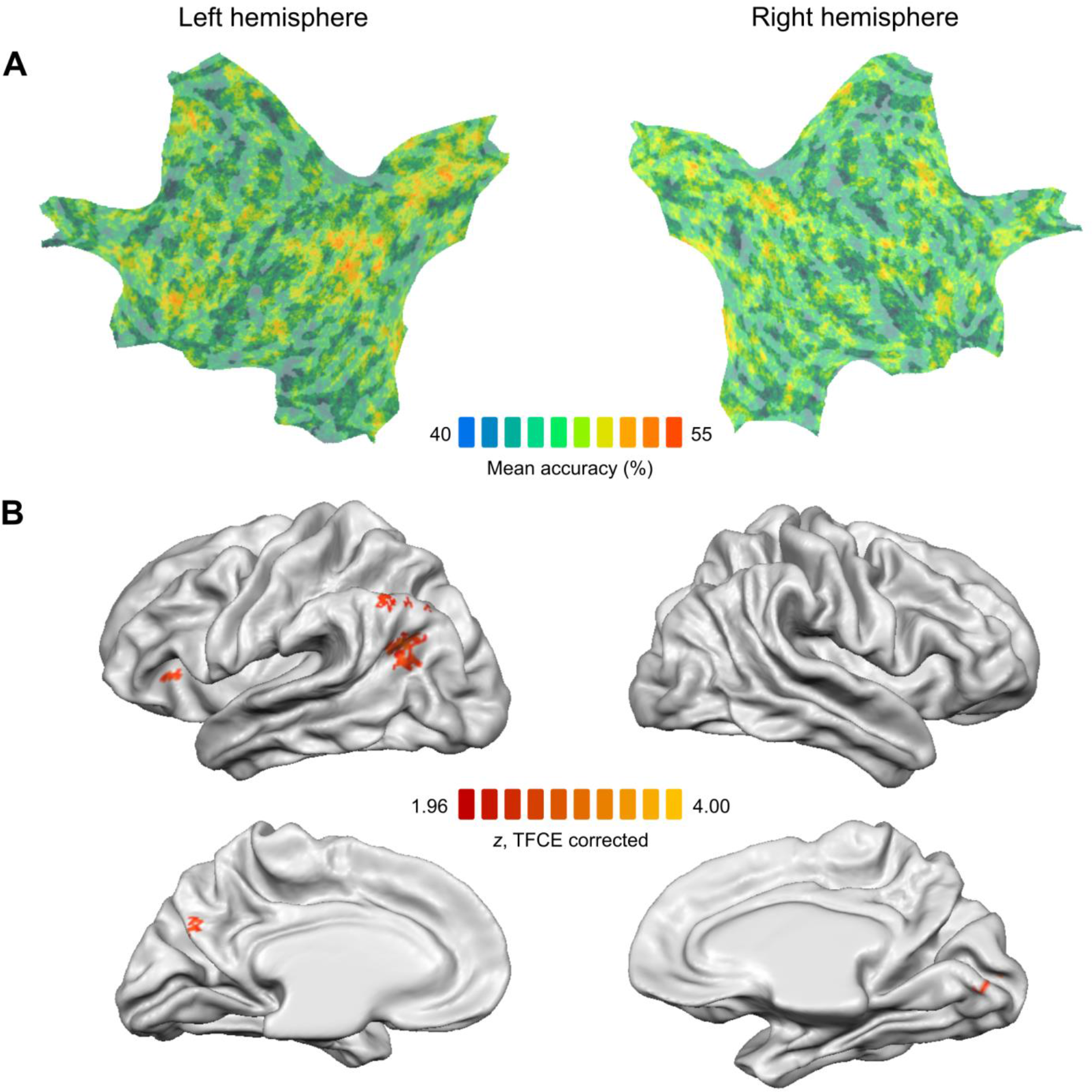
Decoding of trained vs. untrained items, based on the data from the post-training fMRI session. **A.** Mean accuracy maps of the searchlight MVPA. Individual accuracy maps (N = 20) were averaged and projected onto a flattened group-averaged surface. Decoding accuracy at chance is 50%. **B.** Statistical group maps, corrected using TFCE at α = .05 (two-tailed).

## 4. Discussion

We examined the mechanisms underlying naming practice of object and action pictures in healthy participants. We found that, although naming of objects and actions evoked significantly different responses in temporal, parietal and frontal regions, training effects for the two word classes were statistically indistinguishable. Below we will discuss these observations in more detail.

### 4.1. Word class effects

Verbs were produced significantly slower than nouns, in line with the previous reports (Kurland et al., 2018; Vigliocco, Vinson, Lewis, & Garrett, 2004). Converging evidence from univariate and multivariate fMRI analyses points to the bilateral LOTC and predominantly left parietal regions as the potential neural loci of word class effects.

Although the lateral occipital regions surrounding the middle temporal (MT) area are primarily associated with perceptual processing of basic motion (Lingnau, Ashida, Wall, & Smith, 2009; Wall, Lingnau, Ashida, & Smith, 2008), they are also sensitive to motion implied by static images (e.g., Kable, Lease-Spellmeyer, & Chatterjee, 2002; Kourtzi & Kanwisher, 2000), accounting for why these areas may be activated by action drawings. It has recently been proposed that more anterior, temporal regions of the LOTC store conceptual, rather than perceptual, representations of actions (for review, see Lingnau & Downing, 2015). Several findings support this claim. First, stronger activation for verbs than nouns occurs in the lateral posterior temporal regions in response not only to pictures/videos, but also to written and spoken words (Kable, Kan, Wilson, Thompson-Schill, & Chatterjee, 2005; Kable et al., 2002; Papeo & Lingnau, 2015). Second, action-related information in the anterior LOTC does not rely on visual experience, as it is sensitive to both motion verbs and mental state verbs (Bedny, Caramazza, Grossman, Pascual-Leone, & Saxe, 2008; Bedny, Caramazza, Pascual-Leone, & Saxe, 2012), and is activated during action processing in congenitally blind individuals (Bedny et al., 2012). Third, experiments employing MVPA successfully decoded representations of observed actions in portions of the LOTC anterior to MT that generalize across object exemplars and motion kinematics (Wurm, Ariani, Greenlee, & Lingnau, 2016; Wurm & Lingnau, 2015).

Notably, verb-preferring LOTC activations in our experiment extended into the mid-portion of the left middle/superior temporal cortex, a region implicated in retrieval of lexical and grammatical (rather than conceptual/semantic) information about verbs (Crepaldi et al., 2011; Willms et al., 2011). While the conceptual and linguistic accounts cannot be teased apart in the context of our experiment (since verbs referred to actions and nouns referred to objects), studies investigating the nature of verb-related LOTC activations (Bedny, Dravida, & Saxe, 2014; Peelen, Romagno, & Caramazza, 2012) confirm that whereas posterior lateral temporal regions store conceptual representations of actions, a distinct cluster in the left pMTG may be specialized for processing verbs as a grammatical class.

Several clusters in the posterior parietal lobe (on the left in the univariate analysis; bilateral, but mainly on the left in the searchlight analysis) were recruited during action, as compared to object naming. The parietal cortex is the endpoint of the dorsal visual processing stream (Goodale & Milner, 1992; Ungerleider & Mishkin, 1982), mediating spatial perception and visually-guided reaching and grasping movements. In addition to action planning and execution, it is involved in action-related perceptual tasks, such as action observation, and in processing of tools and other action attributes (for review, see Culham & Valyear, 2006). Activation of the posterior parietal cortex has also been reported in language tasks, including verb production (Marangolo, Piras, Galati, & Burani, 2006; Saccuman et al., 2006; Shapiro, Moo, & Caramazza, 2006; Warburton et al., 1996) and comprehension (Tsigka, Papadelis, Braun, & Miceli, 2014). The linguistic nature of parietal activations in our experiment is indirectly supported by their being left-lateralized. While there is no consensus on the role of parietal regions in verb processing, recent studies suggest that they may be crucial for thematic role assignment (Finocchiaro, Capasso, Cattaneo, Zuanazzi, & Miceli, 2015; Thothathiri, Kimberg, & Schwartz, 2012). On an alternative view, increased parietal activations during action naming may be explained by greater complexity of action drawings that for transitive verbs included not only agents, but also undergoers and instruments (Liljeström, Hultén, Parkkonen, & Salmelin, 2009; Liljeström et al., 2008).

In addition, the searchlight analysis revealed that nouns and verbs showed significantly different activation patterns in virtually all left inferior frontal regions engaged in picture naming. The association of verb processing with left IFG activation has a long-standing tradition, corroborated by neuropsychological findings. The fronto-temporal dichotomy hypothesis (FTDH; Damasio & Tranel, 1993) linked selective verb impairment to lesions in the left posterior inferior frontal and precentral gyri, and selective noun retrieval deficits to middle and inferior temporal damage. Consistent evidence was reported in lesion studies (Bak, O’Donovan, Xuereb, Boniface, & Hodges, 2001; Daniele, Giustolisi, Silveri, Colosimo, & Gainotti, 1994; Shapiro & Caramazza, 2003; Tranel, Adolphs, Damasio, & Damasio, 2001; for review, see Cappa & Perani, 2003), cortical mapping in glioma patients (Havas et al., 2015) and neuroimaging experiments (Bedny et al., 2008; Berlingeri et al., 2008; Palti, Ben Shachar, Hendler, & Hadar, 2007; Perani et al., 1999; Tyler, Bright, Fletcher, & Stamatakis, 2004; Tyler et al., 2003). However, evidence from some reports is at odds with the FTDH (e.g., Aggujaro et al., 2006; De Renzi & di Pellegrino, 1995; Silveri & Di Betta, 1997; Silveri, Perri, & Cappa, 2003; Tranel et al., 2008), suggesting that noun/verb dissociations result not from a single factor, but rather from disruption at different levels of word processing, including semantic, phonological and syntactic (Black & Chiat, 2003; Cappa & Perani, 2003). According to an alternative account, activation differences in the verb vs. noun contrast are incidental and are attributable to the difference in morphosyntactic and task demands, rather than the presence of any verb-specific linguistic information in this area (Berlingeri et al., 2008; Siri et al., 2008; Vigliocco et al., 2011). On this view, increased prefrontal response to verbs as compared to nouns in the context of a picture naming task (and during word production in general) may reflect differences in lexical selection demands, as verbs typically have more synonyms/hyponyms/hyperonyms than nouns, and thus place more load on selection processes (Kan & Thompson-Schill, 2004).

Using MVPA, we were able to reliably decode nouns and verbs in posterior ventral occipitotemporal cortices bilaterally, consistent with the role attributed to this area in representation of object concepts (for review, see Martin, 2007). The univariate analysis detected only a small noun-preferring cluster in the left posterior fusiform, which may have to do with the fact that our stimuli were selected from a wide range of semantic categories whose conceptual representations are distributed along the ventral occipitotemporal cortex.

### 4.2. Training effects

Naming latencies were significantly reduced for trained nouns and verbs, both when comparing trained and untrained items in the post-training session and when contrasting the same items from each trained subset across sessions. At the neural level, intensive training for ten days yielded similar activation changes for nouns and verbs, that were separable from the effects of incidental naming practice and involved both anterior and posterior regions of the left hemisphere.

#### 4.2.1. Training-related activation decreases in left anterior regions

Training of nouns and verbs was associated with significant decreases of the BOLD response in the left IFG and anterior insula, replicating findings of studies on repeated object naming (Basso et al., 2013; MacDonald et al., 2015; Meister et al., 2005; Meltzer et al., 2009; van Turennout et al., 2003, 2000). Notably, activation decreases in these regions were significantly greater after the explicit training as compared to a single word repetition over the course of two weeks, attesting to the cumulative nature of practice.

Activation of the left posterior IFG during naming tasks has been ascribed to a number of linguistic functions, including both phonological and semantic processing (Poldrack et al., 1999; Vigneau et al., 2006). Although the practice-related decreases of the BOLD signal in this region may be attributed to facilitation at any level(s) of language processing, they may also be explained by decreased reliance on executive mechanisms, such as response selection and inhibition of competing responses (Thompson-Schill, D’Esposito, Aguirre, & Farah, 1997; Thompson-Schill, D’Esposito, & Kan, 1999). Indeed, subjects were encouraged to settle on target words from the beginning and stick to them throughout the training. This may have increased name agreement, thus decreasing left prefrontal activation (Kan & Thompson-Schill, 2004).

Activation decreases in the left anterior insula support findings of Basso et al. (2013), who compared the BOLD responses to low-frequency nouns before and after training with response to high-frequency nouns that were not involved in practice. While prior to training low-frequency items yielded greater activations in the insula bilaterally, in the post-training session they were indistinguishable from high-frequency items. Hence, training effects in the left anterior insula may mimic the frequency effects, as the usage frequency of trained items was manipulated by intensive repetition. Supporting evidence comes from previous studies implicating insula in processing of low-frequency words (Binder, Medler, Desai, Conant, & Liebenthal, 2005; Carreiras, Mechelli, & Price, 2006; de Zubicaray, McMahon, Eastburn, Finnigan, & Humphreys, 2005; Graves, Grabowski, Mehta, & Gordon, 2007).

#### 4.2.2. Training-related activation changes in left temporal and parietal regions

MVPA distinguished activation patterns associated with naming of trained and untrained words in several posterior regions of the left hemisphere, including the precuneus, angular gyrus and posterior superior temporal sulcus (pSTS).

As discussed above, posterior lateral temporal cortices are implicated in storage of conceptual and lexical representations. Specifically, activation in pSTS is often attributed to processing of lexical word forms (Hickok & Poeppel, 2007; Indefrey & Levelt, 2004). In our study, activation changes in this region might reflect facilitated access to phonological representations of the trained words. Interestingly, sparing of this region was the sole predictor of anomia recovery in a lesion-symptom mapping study (Fridriksson, 2010; Supplementary Fig. 2, blue sphere), attesting to its role in word retrieval. Notably, as shown by diffusion-weighted imaging (DWI) studies in healthy subjects, posterior lateral temporal regions are extensively connected to the eloquent areas in the left IFG by the arcuate fasciculus, both directly and indirectly, via the inferior parietal lobule (Catani, Jones, & ffytche, 2005).

Practice-related increase of the BOLD signal in the precuneus has been reported in several studies of repeated object naming in healthy populations (Basso et al., 2013; Kurland et al., 2018; MacDonald et al., 2015) and in patients with aphasia (Fridriksson, 2010; Fridriksson et al., 2007; Heath et al., 2015). Increased response of the precuneus and the angular gyrus was also reported following sentence repetition (Hasson, Nusbaum, & Small, 2006; Poppenk, McIntosh, & Moscovitch, 2016). Kurland et al. (2018) documented practice effects following several repetitions of nouns and verbs in a portion of the inferior parietal lobule, closely overlapping with the angular gyrus cluster identified by our study (Supplementary Fig. 2, yellow sphere). The above-mentioned studies linked activation of the parietal regions to the explicit memory of practiced items. Evidence consistent with this view was reported by Schott et al. (2005), who found that only conscious recognition of previously studied items (but not priming in the absence of explicit memory) yielded increased response in the precuneus and the inferior parietal lobule.

Whereas the role of the precuneus in mediating episodic memory has been long established, the supporting role of the angular gyrus in this process was highlighted relatively recently (Seghier, 2013; Yazar, Bergström, & Simons, 2012). Importantly, fibers from the angular gyrus project both to the domain-general regions implicated in long-term memory, including the precuneus, and to the IFG (Seghier, 2013), which makes this area well-suited to mediate language learning.

While at first glance training-related changes outside of the classic language circuit may seem surprising, our results map well onto the studies in anomic patients that report a positive relationship between treatment-induced naming improvement and modulation of activity in the lateral and medial parietal cortices (Fridriksson, 2010; Fridriksson et al., 2007; Menke et al., 2009). Although functional changes in parietal regions did not receive much attention in the clinical literature (that mostly focuses on the perisylvian areas and their right-hemispheric homologues), they may support naming recovery in some patients with chronic aphasia. Thus, the intactness and potential functional reorganization of these areas following practice should be considered in naming treatment studies with patients.

## Acknowledgements

The authors thank UNITN students Irene Graziosi and Mariapia Piccinini for assistance with translation and subject recruitment.

## References

Aggujaro, S., Crepaldi, D., Pistarini, C., Taricco, M., & Luzzatti, C. (2006). Neuro-anatomical correlates of impaired retrieval of verbs and nouns: Interaction of grammatical class, imageability and actionality. Journal of Neurolinguistics, 19(3), 175–194. https://doi.org/10.1016/j.jneuroling.2005.07.004

Bak, T. H., O’Donovan, D. G., Xuereb, J. H., Boniface, S., & Hodges, J. R. (2001). Selective impairment of verb processing associated with pathological changes in Brodmann areas 44 and 45 in the motor neurone disease–dementia–aphasia syndrome. Brain, 124(1), 103–120. https://doi.org/10.1093/brain/124.1.103

Basso, G., Magon, S., Reggiani, F., Capasso, R., Monittola, G., Yang, F. J., & Miceli, G. (2013). Distinguishable neurofunctional effects of task practice and item practice in picture naming: A BOLD fMRI study in healthy subjects. Brain and Language, 126(3), 302–313. https://doi.org/10.1016/j.bandl.2013.07.002

Bastiaanse, R., Wieling, M., & Wolthuis, N. (2016). The role of frequency in the retrieval of nouns and verbs in aphasia. Aphasiology, 30(11), 1221–1239. https://doi.org/10.1080/02687038.2015.1100709

Beauchamp, M. S., Petit, L., Ellmore, T. M., Ingeholm, J., & Haxby, J. V. (2001). A parametric fMRI study of overt and covert shifts of visuospatial attention. NeuroImage, 14(2), 310–321. https://doi.org/10.1006/nimg.2001.0788

Bedny, M., Caramazza, A., Grossman, E., Pascual-Leone, A., & Saxe, R. (2008). Concepts are more than percepts: The case of action verbs. The Journal of Neuroscience, 28(44), 11347–11353. https://doi.org/10.1523/jneurosci.3039-08.2008

Bedny, M., Caramazza, A., Pascual-Leone, A., & Saxe, R. (2012). Typical neural representations of action verbs develop without vision. Cerebral Cortex, 22(2), 286–293. https://doi.org/10.1093/cercor/bhr081

Bedny, M., Dravida, S., & Saxe, R. (2014). Shindigs, brunches, and rodeos: The neural basis of event words. Cognitive, Affective, and Behavioral Neuroscience, 14(3), 891–901. https://doi.org/10.3758/s13415-013-0217-z

Benjamini, Y., & Yekutieli, D. (2001). The control of the false discovery rate in multiple testing under dependency. Annals of Statistics, 29(4), 1165–1188. https://doi.org/10.1214/aos/1013699998

Berlingeri, M., Crepaldi, D., Roberti, R., Scialfa, G., Luzzatti, C., & Paulesu, E. (2008). Nouns and verbs in the brain: Grammatical class and task specific effects as revealed by fMRI. Cognitive Neuropsychology, 25(4), 528–558. https://doi.org/10.1080/02643290701674943

Bertinetto, P. M., Burani, C., Laudanna, A., Marconi, L., Ratti, D., Rolando, C., & Thornton, A. M. (2005). Corpus e Lessico di Frequenza dell’Italiano Scritto (CoLFIS). Retrieved from http://linguistica.sns.it/CoLFIS/Home.htm

Binder, J. R., Medler, D. A., Desai, R. H., Conant, L. L., & Liebenthal, E. (2005). Some neurophysiological constraints on models of word naming. NeuroImage, 27(3), 677–693. https://doi.org/10.1016/j.neuroimage.2005.04.029

Black, M., & Chiat, S. (2003). Noun–verb dissociations: A multi-faceted phenomenon. Journal of Neurolinguistics, 16(2–3), 231–250. https://doi.org/10.1016/s0911-6044(02)00017-9

Brainard, D. H. (1997). The Psychophysics Toolbox. Spatial Vision, 10, 433–436. https://doi.org/10.1163/156856897x00357

Cappa, S. F., & Perani, D. (2003). The neural correlates of noun and verb processing. Journal of Neurolinguistics, 16(2–3), 183–189. https://doi.org/10.1016/s0911-6044(02)00013-1

Carreiras, M., Mechelli, A., & Price, C. J. (2006). Effect of word and syllable frequency on activation during lexical decision and reading aloud. Human Brain Mapping, 27(12), 963–972. https://doi.org/10.1002/hbm.20236

Catani, M., Jones, D. K., & ffytche, D. H. (2005). Perisylvian language networks of the human brain. Annals of Neurology, 57(1), 8–16. https://doi.org/10.1002/ana.20319

Corbetta, M., & Shulman, G. L. (1998). Human cortical mechanisms of visual attention during orienting and search. Philosophical Transactions of the Royal Society B: Biological Sciences, 353(1373), 1353–1362. https://doi.org/10.1098/rstb.1998.0289

Cousineau, D. (2005). Confidence intervals in within-subject designs: A simpler solution to Loftus and Masson’s method. Tutorials in Quantitative Methods for Psychology, 1(1), 42–45. https://doi.org/10.20982/tqmp.01.1.p042

Crepaldi, D., Berlingeri, M., Paulesu, E., & Luzzatti, C. (2011). A place for nouns and a place for verbs? A critical review of neurocognitive data on grammatical-class effects. Brain and Language, 116(1), 33–49. https://doi.org/10.1016/j.bandl.2010.09.005

Culham, J. C., & Valyear, K. F. (2006). Human parietal cortex in action. Current Opinion in Neurobiology, 16(2), 205–212. https://doi.org/10.1016/j.conb.2006.03.005

Damasio, A. R., & Tranel, D. (1993). Nouns and verbs are retrieved with differently distributed neural systems. Proceedings of the National Academy of Sciences of the USA, 90(11), 4957–4960. https://doi.org/10.1073/pnas.90.11.4957

Daniele, A., Giustolisi, L., Silveri, M. C., Colosimo, C., & Gainotti, G. (1994). Evidence for a possible neuroanatomical basis for lexical processing of nouns and verbs. Neuropsychologia, 32(11), 1325–1341. https://doi.org/10.1016/0028-3932(94)00066-2

Davis, T., LaRocque, K. F., Mumford, J. A., Norman, K. A., Wagner, A. D., & Poldrack, R. A. (2014). What do differences between multi-voxel and univariate analysis mean? How subject-, voxel-, and trial-level variance impact fMRI analysis. NeuroImage, 97, 271–283. https://doi.org/10.1016/j.neuroimage.2014.04.037

De Renzi, E., & di Pellegrino, G. (1995). Sparing of verbs and preserved, but ineffectual reading in a patient with impaired word production. Cortex, 31(4), 619–636. https://doi.org/10.1016/s0010-9452(13)80016-0

de Zubicaray, G. I., McMahon, K. L., Eastburn, M. M., Finnigan, S., & Humphreys, M. S. (2005). fMRI evidence of word frequency and strength effects during episodic memory encoding. Cognitive Brain Research, 22(3), 439–450. https://doi.org/10.1016/j.cogbrainres.2004.10.002

Etzel, J. A., Zacks, J. M., & Braver, T. S. (2013). Searchlight analysis: Promise, pitfalls, and potential. NeuroImage, 78, 261–269. https://doi.org/10.1016/j.neuroimage.2013.03.041

Finocchiaro, C., Capasso, R., Cattaneo, L., Zuanazzi, A., & Miceli, G. (2015). Thematic role assignment in the posterior parietal cortex: A TMS study. Neuropsychologia, 77, 223–232. https://doi.org/10.1016/j.neuropsychologia.2015.08.025

Fischl, B., Sereno, M. I., Tootell, R. B. H., & Dale, A. M. (1999). High-resolution intersubject averaging and a coordinate system for the cortical surface. Human Brain Mapping, 8(4), 272–284. https://doi.org/10.1002/(SICI)1097-0193(1999)8

Forsythe, A., Mulhern, G., & Sawey, M. (2008). Confounds in pictorial sets: The role of complexity and familiarity in basic-level picture processing. Behavior Research Methods, 40(1), 116–129. https://doi.org/10.3758/brm.40.1.116

Fridriksson, J. (2010). Preservation and modulation of specific left hemisphere regions is vital for treated recovery from anomia in stroke. The Journal of Neuroscience, 30(35), 11558–11564. https://doi.org/10.1523/jneurosci.2227-10.2010

Fridriksson, J., Moser, D., Bonilha, L., Morrow-Odom, K. L., Shaw, H., Fridriksson, A., … Rorden, C. (2007). Neural correlates of phonological and semantic-based anomia treatment in aphasia. Neuropsychologia, 45(8), 1812–1822. https://doi.org/10.1016/j.neuropsychologia.2006.12.017

Friston, K. J., Fletcher, P., Josephs, O., Holmes, A., Rugg, M. D., & Turner, R. (1998). Event-related fMRI: Characterizing differential responses. NeuroImage, 7(1), 30–40. https://doi.org/10.1006/nimg.1997.0306

Goebel, R., Esposito, F., & Formisano, E. (2006). Analysis of Functional Image Analysis Contest (FIAC) data with BrainVoyager QX: From single-subject to cortically aligned group General Linear Model analysis and self-organizing group Independent Component Analysis. Human Brain Mapping, 27(5), 392–401. https://doi.org/10.1002/hbm.20249

Goodale, M. A., & Milner, A. D. (1992). Separate visual pathways for perception and action. Trends in Neurosciences, 15(1), 20–25. https://doi.org/10.1016/0166-2236(92)90344-8

Goodale, M. A., & Milner, A. D. (2018). Two visual pathways – Where have they taken us and where will they lead in future? Cortex, 98, 283–292. https://doi.org/10.1016/j.cortex.2017.12.002

Graves, W. W., Grabowski, T. J., Mehta, S., & Gordon, J. K. (2007). A neural signature of phonological access: Distinguishing the effects of word frequency from familiarity and length in overt picture naming. Journal of Cognitive Neuroscience, 19(4), 617–631. https://doi.org/10.1162/jocn.2007.19.4.617

Hasson, U., Nusbaum, H. C., & Small, S. L. (2006). Repetition suppression for spoken sentences and the effect of task demands. Journal of Cognitive Neuroscience, 18(12), 2013–2029. https://doi.org/10.1162/jocn.2006.18.12.2013

Havas, V., Gabarrós, A., Juncadella, M., Rifa-Ros, X., Plans, G., Acebes, J. J., … Rodríguez-Fornells, A. (2015). Electrical stimulation mapping of nouns and verbs in Broca’s area. Brain and Language, 145–146, 53–63. https://doi.org/10.1016/j.bandl.2015.04.005

Haxby, J. V. (2012). Multivariate pattern analysis of fMRI: The early beginnings. NeuroImage, 62(2), 852–855. https://doi.org/10.1016/j.neuroimage.2012.03.016

Haxby, J. V., Gobbini, M. I., Furey, M. L., Ishai, A., Schouten, J. L., & Pietrini, P. (2001). Distrubuted and overlapping representations of face and objects in ventral temporal cortex. Science, 293(5539), 2425–2430. https://doi.org/10.1126/science.1063736

Heath, S., McMahon, K. L., Nickels, L. A., Angwin, A. J., MacDonald, A. D., van Hees, S., … Copland, D. A. (2015). An fMRI investigation of the effects of attempted naming on word retrieval in aphasia. Frontiers in Human Neuroscience, 9, 291. https://doi.org/10.3389/fnhum.2015.00291

Henson, R. N. A. (2003). Neuroimaging studies of priming. Progress in Neurobiology, 70(1), 53–81. https://doi.org/10.1016/S0301-0082(03)00086-8

Hickok, G., & Poeppel, D. (2007). The cortical organisation of speech processing. Nature Reviews Neuroscience, 8(5), 393–402. https://doi.org/10.1038/nrn2113

Howard, D. (2000). Cognitive neuropsychology and aphasia therapy: The case of word retrieval. In I. Papathanasiou (Ed.), Acquired neurogenic communication disorders: A clinical perspective (pp. 76–99). London: Whurr.

Indefrey, P., & Levelt, W. J. M. (2004). The spatial and temporal signatures of word production components. Cognition, 92(1–2), 101–144. https://doi.org/10.1016/j.cognition.2002.06.001

Jimura, K., & Poldrack, R. A. (2012). Analyses of regional-average activation and multivoxel pattern information tell complementary stories. Neuropsychologia, 50(4), 544–552. https://doi.org/10.1016/j.neuropsychologia.2011.11.007

Kable, J. W., Kan, I. P., Wilson, A., Thompson-Schill, S. L., & Chatterjee, A. (2005). Conceptual representations of action in the lateral temporal cortex. Journal of Cognitive Neuroscience, 17(12), 1855–1870. https://doi.org/10.1162/089892905775008625

Kable, J. W., Lease-Spellmeyer, J., & Chatterjee, A. (2002). Neural substrates of action event knowledge. Journal of Cognitive Neuroscience, 14(5), 795–805. https://doi.org/10.1162/08989290260138681

Kan, I. P., & Thompson-Schill, S. L. (2004). Effect of name agreement on prefrontal activity during overt and covert picture naming. Cognitive, Affective, and Behavioral Neuroscience, 4(1), 43–57. https://doi.org/10.3758/cabn.4.1.43

Kourtzi, Z., & Kanwisher, N. (2000). Activation in human MT/MST by static images with implied motion. Journal of Cognitive Neuroscience, 12(1), 48–55. https://doi.org/10.1162/08989290051137594

Kriegeskorte, N., Goebel, R., & Bandettini, P. (2006). Information-based functional brain mapping. Proceedings of the National Academy of Sciences of the USA, 103(10), 3863–3868. https://doi.org/10.1073/pnas.0600244103

Kurland, J., Liu, A., & Stokes, P. (2018). Practice effects in healthy older adults: Implications for treatment-induced neuroplasticity in Aphasia. Neuropsychologia, 109, 116–125. https://doi.org/10.1016/j.neuropsychologia.2017.12.003

Liljeström, M., Hultén, A., Parkkonen, L., & Salmelin, R. (2009). Comparing MEG and fMRI views to naming actions and objects. Human Brain Mapping, 30(6), 1845–1856. https://doi.org/10.1002/hbm.20785

Liljeström, M., Tarkiainen, A., Parviainen, T., Kujala, J., Numminen, J., Hiltunen, J., … Salmelin, R. (2008). Perceiving and naming actions and objects. NeuroImage, 41(3), 1132–1141. https://doi.org/10.1016/j.neuroimage.2008.03.016

Lingnau, A., Ashida, H., Wall, M. B., & Smith, A. T. (2009). Speed encoding in human visual cortex revealed by fMRI adaptation. Journal of Vision, 9(13), 1–14. https://doi.org/10.1167/9.13.3

Lingnau, A., & Downing, P. E. (2015). The lateral occipitotemporal cortex in action. Trends in Cognitive Sciences, 19(5), 268–277. https://doi.org/10.1016/j.tics.2015.03.006

MacDonald, A. D., Heath, S., McMahon, K. L., Nickels, L. A., Angwin, A. J., van Hees, S., … Copland, D. A. (2015). Neuroimaging the short- and long-term effects of repeated picture naming in healthy older adults. Neuropsychologia, 75, 170–178. https://doi.org/10.1016/j.neuropsychologia.2015.06.007

Marangolo, P., Piras, F., Galati, G., & Burani, C. (2006). Functional anatomy of derivational morphology. Cortex, 42(8), 1093–1106. https://doi.org/10.1016/S0010-9452(08)70221-1

Martin, A. (2007). The representation of object concepts in the brain. Annual Review of Psychology, 58(1), 25–45. https://doi.org/10.1146/annurev.psych.57.102904.190143

Mätzig, S., Druks, J., Masterson, J., & Vigliocco, G. (2009). Noun and verb differences in picture naming: Past studies and new evidence. Cortex, 45(6), 738–758. https://doi.org/10.1016/j.cortex.2008.10.003

Meister, I. G., Weidemann, J., Foltys, H., Brand, H., Willmes, K., Krings, T., … Boroojerdi, B. (2005). The neural correlate of very-long-term picture priming. European Journal of Neuroscience, 21(4), 1101–1106. https://doi.org/10.1111/j.1460-9568.2005.03941.x

Meltzer, J. A., Postman-Caucheteux, W. A., McArdle, J. J., & Braun, A. R. (2009). Strategies for longitudinal neuroimaging studies of overt language production. NeuroImage, 47(2), 745–755. https://doi.org/10.1016/j.neuroimage.2009.04.089

Menke, R., Meinzer, M., Kugel, H., Deppe, M., Baumgärtner, A., Schiffbauer, H., … Breitenstein, (2009). Imaging short- and long-term training success in chronic aphasia. BMC Neuroscience, 10(1), 118. https://doi.org/10.1186/1471-2202-10-118

Miceli, G., Laudanna, A., & Capasso, R. (2001). Batteria per l’analisi dei deficit afasici. Bologna: ECM.

Nichols, T. E., & Holmes, A. P. (2002). Nonparametric permutation tests for functional neuroimaging: A primer with examples. Human Brain Mapping, 15(1), 1–25. https://doi.org/10.1002/hbm.1058

Nickels, L. A. (2002). Improving word finding: Practice makes (closer to) perfect? Aphasiology, 16(10–11), 1047–1060. https://doi.org/10.1080/02687040143000618

Nobre, A. C., Sebestyen, G. N., Gitelman, D. R., Mesulam, M. M., Frackowiak, R. S. J., & Frith, C. (1997). Functional localization of the system for visuospatial attention using positron emission tomography. Brain, 120(3), 515–533. https://doi.org/10.1093/brain/120.3.515

Oldfield, R. C. (1971). The assessment and analysis of handedness: The Edinburgh inventory. Neuropsychologia, 9(1), 97–113. https://doi.org/10.1016/0028-3932(71)90067-4

Oosterhof, N. N., Connolly, A. C., & Haxby, J. V. (2016). CoSMoMVPA: Multi-modal multivariate pattern analysis of neuroimaging data in Matlab/GNU Octave. Frontiers in Neuroinformatics, 10, 1–27. https://doi.org/10.3389/fninf.2016.00027

Oosterhof, N. N., Wiestler, T., Downing, P. E., & Diedrichsen, J. (2011). A comparison of volume-based and surface-based multi-voxel pattern analysis. NeuroImage, 56(2), 593–600. https://doi.org/10.1016/j.neuroimage.2010.04.270

Palti, D., Ben Shachar, M., Hendler, T., & Hadar, U. (2007). Neural correlates of semantic and morphological processing of Hebrew nouns and verbs. Human Brain Mapping, 28(4), 303–314. https://doi.org/10.1002/hbm.20280

Papeo, L., & Lingnau, A. (2015). First-person and third-person verbs in visual motion-perception regions. Brain and Language, 141, 135–141. https://doi.org/10.1016/j.bandl.2014.11.011

Peelen, M. V., Romagno, D., & Caramazza, A. (2012). Independent representations of verbs and actions in left lateral temporal cortex. Journal of Cognitive Neuroscience, 24(10), 2096–2107. https://doi.org/10.1162/jocn_a_00257

Perani, D., Cappa, S. F., Schnur, T., Tettamanti, M., Collina, S., Rosa, M. M., & Fazio, F. (1999). The neural correlates of verb and noun processing: A PET study. Brain, 122(12), 2337–2344. https://doi.org/10.1093/brain/122.12.2337

Poldrack, R. A., Wagner, A. D., Prull, M. W., Desmond, J. E., Glover, G. H., & Gabrieli, J. D. E. (1999). Functional specialization for semantic and phonological processing in the left inferior prefrontal cortex. NeuroImage, 10(1), 15–35. https://doi.org/10.1006/nimg.1999.0441

Poppenk, J., McIntosh, A. R., & Moscovitch, M. (2016). fMRI evidence of equivalent neural suppression by repetition and prior knowledge. Neuropsychologia, 90, 159–169. https://doi.org/10.1016/j.neuropsychologia.2016.06.034

Saccuman, M. C., Cappa, S. F., Bates, E. A., Arevalo, A., Della Rosa, P., Danna, M., & Perani, D. (2006). The impact of semantic reference on word class: An fMRI study of action and object naming. NeuroImage, 32(4), 1865–1878. https://doi.org/10.1016/j.neuroimage.2006.04.179

Schacter, D. L., & Buckner, R. L. (1998). Priming and the brain. Neuron, 20(2), 185–195. https://doi.org/10.1016/s0896-6273(00)80448-1

Schott, B. H., Henson, R. N. A., Richardson-Klavehn, A., Becker, C., Thoma, V., Heinze, H.-J., & Düzel, E. (2005). Redefining implicit and explicit memory: The functional neuroanatomy of priming, remembering, and control of retrieval. Proceedings of the National Academy of Sciences of the USA, 102(4), 1257–1262. https://doi.org/10.1073/pnas.0409070102

Schwarzbach, J. (2011). A simple framework (ASF) for behavioral and neuroimaging experiments based on the psychophysics toolbox for MATLAB. Behavior Research Methods, 43(4), 1194–1201. https://doi.org/10.3758/s13428-011-0106-8

Seghier, M. L. (2013). The angular gyrus: Multiple functions and multiple subdivisions. Neuroscientist, 19(1), 43–61. https://doi.org/10.1177/1073858412440596

Shapiro, K. A., & Caramazza, A. (2003). Grammatical processing of nouns and verbs in left frontal cortex? Neuropsychologia, 41(9), 1189–1198. https://doi.org/10.1016/s0028-3932(03)00037-x

Shapiro, K. A., Moo, L. R., & Caramazza, A. (2006). Cortical signatures of noun and verb production. Proceedings of the National Academy of Sciences of the USA, 103(5), 1644–1649. https://doi.org/10.1073/pnas.0504142103

Silveri, M. C., & Di Betta, A. M. (1997). Noun-verb dissociations in brain-damaged patients: Further evidence. Neurocase, 3(6), 477–488. https://doi.org/10.1093/neucas/3.6.477

Silveri, M. C., Perri, R., & Cappa, A. (2003). Grammatical class effects in brain-damaged patients: Functional locus of noun and verb deficit. Brain and Language, 85(1), 49–66. https://doi.org/10.1016/s0093-934x(02)00504-7

Siri, S., Tettamanti, M., Cappa, S. F., Della Rosa, P., Saccuman, C., Scifo, P., & Vigliocco, G. (2008). The neural substrate of naming events: Effects of processing demands but not of grammatical class. Cerebral Cortex, 18(1), 171–177. https://doi.org/10.1093/cercor/bhm043

Smith, S. M., & Nichols, T. E. (2009). Threshold-free cluster enhancement: Addressing problems of smoothing, threshold dependence and localisation in cluster inference. NeuroImage, 44(1), 83–98. https://doi.org/10.1016/j.neuroimage.2008.03.061

Talairach, J., & Tournoux, P. (1988). Co-planar stereotaxic atlas of the human brain: 3-dimensional proportional system: An approach to cerebral imaging. New York, NY: Thieme Medical Publishers.

Thompson-Schill, S. L., D’Esposito, M., Aguirre, G. K., & Farah, M. J. (1997). Role of left inferior prefrontal cortex in retrieval of semantic knowledge: A reevaluation. Proceedings of the National Academy of Sciences of the USA, 94(26), 14792–14797. https://doi.org/10.1073/pnas.94.26.14792

Thompson-Schill, S. L., D’Esposito, M., & Kan, I. P. (1999). Effects of repetition and competition on activity in left prefrontal cortex during word generation. Neuron, 23(3), 513–522. https://doi.org/10.1016/s0896-6273(00)80804-1

Thothathiri, M., Kimberg, D. Y., & Schwartz, M. F. (2012). The neural basis of reversible sentence comprehension: Evidence from voxel-based lesion symptom mapping in aphasia. Journal of Cognitive Neuroscience, 24(1), 212–222. https://doi.org/10.1162/jocn_a_00118

Tranel, D., Adolphs, R., Damasio, H., & Damasio, A. R. (2001). A neural basis for the retrieval of words for actions. Cognitive Neuropsychology, 18(7), 655–674. https://doi.org/10.1080/02643290143000015

Tranel, D., Manzel, K., Asp, E., & Kemmerer, D. (2008). Naming dynamic and static actions: Neuropsychological evidence. Journal of Physiology – Paris, 102(1–3), 80–94. https://doi.org/10.1016/j.jphysparis.2008.03.008

Tsigka, S., Papadelis, C., Braun, C., & Miceli, G. (2014). Distinguishable neural correlates of verbs and nouns: A MEG study on homonyms. Neuropsychologia, 54, 87–97. https://doi.org/10.1016/j.neuropsychologia.2013.12.018

Tyler, L. K., Bright, P., Fletcher, P., & Stamatakis, E. A. (2004). Neural processing of nouns and verbs: The role of inflectional morphology. Neuropsychologia, 42(4), 512–523. https://doi.org/10.1016/j.neuropsychologia.2003.10.001

Tyler, L. K., Stamatakis, E. A., Dick, E., Bright, P., Fletcher, P., & Moss, H. (2003). Objects and their actions: Evidence for a neurally distributed semantic system. NeuroImage, 18(2), 542–557. https://doi.org/10.1016/S1053-8119(02)00047-2

Ungerleider, L. G., & Mishkin, M. (1982). Two cortical visual systems. In D. J. Ingle, M. A. Goodale, & R. J. W. Mansfield (Eds.), Analysis of visual behavior (pp. 549–586). Cambridge, MA: MIT Press.

van Turennout, M., Bielamowicz, L., & Martin, A. (2003). Modulation of neural activity during object naming: Effects of time and practice. Cerebral Cortex, 13(4), 381–391. https://doi.org/10.1093/cercor/13.4.381

van Turennout, M., Ellmore, T., & Martin, A. (2000). Long-lasting cortical plasticity in the object naming system. Nature Neuroscience, 3(12), 1329–1334. https://doi.org/10.1038/81873

Vigliocco, G., Vinson, D. P., Druks, J., Barber, H., & Cappa, S. F. (2011). Nouns and verbs in the brain: A review of behavioural, electrophysiological, neuropsychological and imaging studies. Neuroscience and Biobehavioral Reviews, 35(3), 407–426. https://doi.org/10.1016/j.neubiorev.2010.04.007

Vigliocco, G., Vinson, D. P., Lewis, W., & Garrett, M. F. (2004). Representing the meanings of object and action words: The featural and unitary semantic space hypothesis. Cognitive Psychology, 48(4), 422–488. https://doi.org/10.1016/j.cogpsych.2003.09.001

Vigneau, M., Beaucousin, V., Hervé, P. Y., Duffau, H., Crivello, F., Houdé, O., … Tzourio-Mazoyer, N. (2006). Meta-analyzing left hemisphere language areas: Phonology, semantics, and sentence processing. NeuroImage, 30(4), 1414–1432. https://doi.org/10.1016/j.neuroimage.2005.11.002

Wall, M. B., Lingnau, A., Ashida, H., & Smith, A. T. (2008). Selective visual responses to expansion and rotation in the human MT complex revealed by functional magnetic resonance imaging adaptation. European Journal of Neuroscience, 27(10), 2747–2757. https://doi.org/10.1111/j.1460-9568.2008.06249.x

Warburton, E., Wise, R. J. S., Price, C. J., Weiller, C., Hadar, U., Ramsay, S., & Frackowiak, R. S. J. (1996). Noun and verb retrieval by normal subjects: Studies with PET. Brain, 119(1), 159–179. https://doi.org/10.1093/brain/119.1.159

Willms, J. L., Shapiro, K. A., Peelen, M. V., Pajtas, P. E., Costa, A., Moo, L. R., & Caramazza, A. (2011). Language-invariant verb processing regions in Spanish-English bilinguals. NeuroImage, 57(1), 251–261. https://doi.org/10.1016/j.neuroimage.2011.04.021

Wurm, M. F., Ariani, G., Greenlee, M. W., & Lingnau, A. (2016). Decoding concrete and abstract action representations during explicit and implicit conceptual processing. Cerebral Cortex, 26(8), 3390–3401. https://doi.org/10.1093/cercor/bhv169

Wurm, M. F., & Lingnau, A. (2015). Decoding actions at different levels of abstraction. The Journal of Neuroscience, 35(20), 7727–7735. https://doi.org/10.1523/jneurosci.0188-15.2015

Yazar, Y., Bergström, Z. M., & Simons, J. S. (2012). What is the parietal lobe contribution to long-term memory? Cortex, 48(10), 1381–1382. https://doi.org/10.1016/j.cortex.2012.05.011

Zaitsev, M., Hennig, J., & Speck, O. (2004). Point spread function mapping with parallel imaging techniques and high acceleration factors: Fast, robust, and flexible method for echo-planar imaging distortion correction. Magnetic Resonance in Medicine, 52(5), 1156–1166. https://doi.org/10.1002/mrm.20261

Zeng, H., & Constable, R. T. (2002). Image distortion correction in EPI: Comparison of field mapping with point spread function mapping. Magnetic Resonance in Medicine, 48(1), 137–146. https://doi.org/10.1002/mrm.10200

