## Supplementary Materials for "Neural correlates of object and action naming practice"

Supplementary Figures

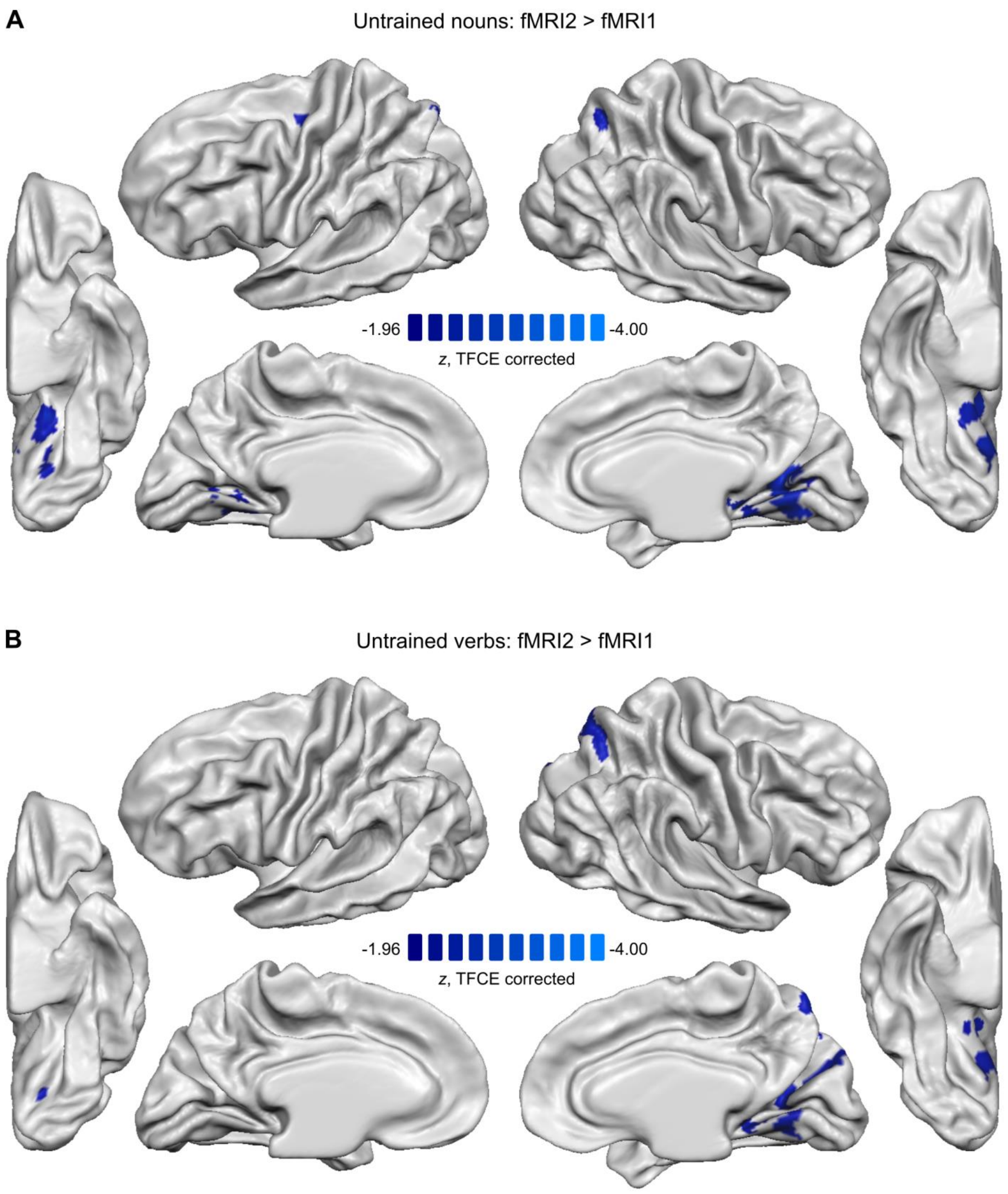


**Supplementary Fig. 1** – Session effects, revealed by the univariate analysis. Effects of stimulus/task habituation, identified by contrasting untrained nouns (***A***) and untrained verbs (***B***) across the two fMRI sessions. The statistical group map (N = 20) for each hemisphere was corrected for multiple comparisons using TFCE ($\alpha$ = .05, two-tailed) and projected onto the group-averaged surface meshes for visualization.

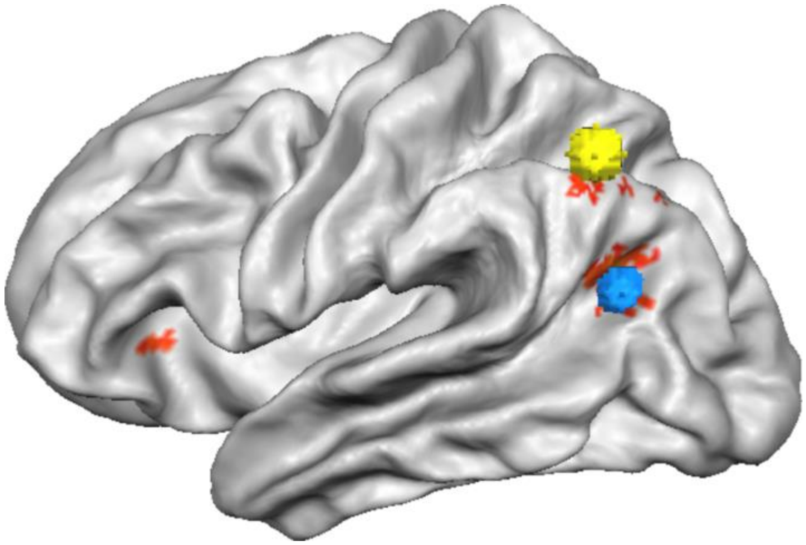

**Supplementary Fig. 2** – Overlapping results of the multivariate analysis of training effects in our study (red clusters) with the areas identified in a lesion-mapping study of anomia treatment (Fridriksson, 2010; blue sphere, posterior temporal lobe, $xyz_{TAL}$ = -36, -64, 18) and a naming practice study with healthy individuals (Kurland et al., 2018; yellow sphere, angular gyrus, $xyz_{TAL}$ = -42, -59, 43). Spheres were created with a 5-mm radius around the coordinates reported in the corresponding studies. The coordinates were converted from the MNI to the Talairach space using the Yale BioImage Suite Package (http://www.bioimagesuite.org).

### Supplementary Tables

**Supplementary Table 1. Active clusters identified in whole-brain univariate analyses.** To obtain cluster statistics, we projected the surface maps back into the volume. The table shows average *z*-values and extent (in $mm^3$) of active clusters, as well as the Talairach coordinates of each cluster's center-of-gravity (COG), its corresponding *z*-value and anatomical label. For clusters larger than 5000 $mm^3$ local maxima were identified using the NeuroElf function *clustervol.m*. Clusters with volume smaller than 10 $mm^3$ are not reported. Two maps marked by an asterisk (*) were thresholded at $z > 1.65$ (one-tailed). The remaining maps were thresholded at $z > 1.96$ (two-tailed). LH = left hemisphere, RH = right hemisphere.

| **cluster** | | | **local maxima** | | | | |
|---|---|---|---|---|---|---|---|
| **side** | **mean *Z*** | **$mm^3$** | ***Z*** | $x_{COG}$ | $y_{COG}$ | $z_{COG}$ | **anatomical region** |
| *Object naming: objects > scrambles in fMRI1 (S1_NU + S1_NT > S1_Control)* | | | | | | | |
| LH | 2.87 | 29590 | 3.09 | -21 | 15 | -10 | Orbital gyrus |
| | | | 3.09 | -23 | -6 | -14 | Parahippocampal gyrus |
| | | | 3.09 | -25 | -40 | -10 | Fusiform gyrus |
| | | | 3.09 | -28 | 27 | -3 | Orbital gyrus |
| | | | 3.09 | -27 | -76 | 22 | Superior parietal lobule |
| | | | 3.09 | -35 | -39 | -15 | Fusiform gyrus |
| | | | 3.09 | -30 | -4 | -24 | Parahippocampal gyrus |
| | | | 3.09 | -39 | 21 | 12 | Inferior frontal gyrus (frontal operculum) |
| | | | 3.09 | -32 | -80 | -13 | Inferior occipital gyrus |
| | | | 3.09 | -34 | -80 | 8 | Middle occipital gyrus |
| | | | 3.09 | -37 | -19 | -19 | Fusiform gyrus |
| | | | 3.09 | -39 | -64 | -9 | Fusiform gyrus |
| | | | 3.09 | -38 | 5 | 32 | Inferior frontal gyrus (pars opercularis) |
| | | | 3.09 | -39 | -73 | 20 | Middle occipital gyrus |
| | | | 3.09 | -42 | -72 | 0 | Inferior occipital gyrus |
| | | | 3.09 | -49 | -56 | -6 | Middle temporal gyrus |
| LH | 2.22 | 683 | 2.46 | -7 | -53 | 10 | Precuneus |
| LH | 2.09 | 719 | 2.17 | -5 | 20 | 48 | Superior frontal gyrus (pre-SMA) |
| RH | 2.72 | 16800 | 3.09 | 43 | -63 | -4 | Inferior occipital gyrus |
| | | | 3.09 | 38 | -60 | -11 | Fusiform gyrus |
| | | | 3.09 | 35 | -75 | -10 | Inferior occipital gyrus |
| | | | 3.09 | 36 | -77 | 10 | Middle occipital gyrus |
| | | | 3.09 | 32 | -80 | 4 | Middle occipital gyrus |
| | | | 3.09 | 38 | -40 | -15 | Fusiform gyrus |
| | | | 3.09 | 28 | -41 | -12 | Fusiform gyrus |
| | | | 3.09 | 27 | -86 | -1 | Middle occipital gyrus |
| | | | 3.09 | 34 | -19 | -18 | Parahippocampal gyrus |
| | | | 3.09 | 28 | -33 | -17 | Fusiform gyrus |
| | | | 2.65 | 28 | -2 | -20 | Parahippocampal gyrus |

| | | | | | | | |
|---|---|---|---|---|---|---|---|
| | | | 2.51 | 27 | 25 | -5 | Orbital gyrus |
| RH | 2.08 | 542 | 2.17 | 22 | -74 | 30 | Superior parietal lobule |
| *Action naming: actions > scrambles in fMRI1 (S1_VU + S1_VT > S1_Control)* | | | | | | | |
| LH | 2.91 | 31331 | 3.09 | -21 | -74 | 33 | Superior parietal lobule |
| | | | 3.09 | -26 | -77 | 23 | Superior parietal lobule |
| | | | 3.09 | -24 | 14 | -9 | Anterior insula |
| | | | 3.09 | -24 | -6 | -17 | Parahippocampal gyrus |
| | | | 3.09 | -25 | -36 | -11 | Fusiform gyrus |
| | | | 3.09 | -29 | 25 | -3 | Orbital gyrus |
| | | | 3.09 | -31 | -21 | -19 | Parahippocampal gyrus |
| | | | 3.09 | -29 | -86 | -2 | Middle occipital gyrus |
| | | | 3.09 | -34 | -78 | 21 | Middle occipital gyrus |
| | | | 3.09 | -37 | -38 | -15 | Fusiform gyrus |
| | | | 3.09 | -39 | 23 | 16 | Inferior frontal gyrus (frontal operculum) |
| | | | 3.09 | -41 | -59 | -9 | Fusiform gyrus |
| | | | 3.09 | -38 | 3 | 32 | Inferior frontal gyrus (pars opercularis) |
| | | | 3.09 | -40 | -69 | 13 | Middle temporal gyrus |
| | | | 3.09 | -41 | -74 | -2 | Inferior occipital gyrus |
| | | | 3.09 | -46 | 15 | 19 | Inferior frontal gyrus (pars triangularis) |
| | | | 3.09 | -50 | -56 | 3 | Middle temporal gyrus |
| LH | 2.25 | 711 | 2.51 | -6 | 20 | 48 | Superior frontal gyrus (pre-SMA) |
| LH | 2.20 | 565 | 2.37 | -7 | -53 | 10 | Precuneus |
| RH | 2.86 | 22670 | 3.09 | 43 | -63 | 4 | Middle temporal gyrus |
| | | | 3.09 | 39 | -75 | 10 | Middle occipital gyrus |
| | | | 3.09 | 39 | -60 | -11 | Fusiform gyrus |
| | | | 3.09 | 42 | -46 | -12 | Fusiform gyrus |
| | | | 3.09 | 35 | -74 | -11 | Inferior occipital gyrus |
| | | | 3.09 | 39 | -15 | -21 | Fusiform gyrus |
| | | | 3.09 | 32 | -79 | 5 | Middle occipital gyrus |
| | | | 3.09 | 34 | -36 | -16 | Fusiform gyrus |
| | | | 3.09 | 30 | -44 | -13 | Fusiform gyrus |
| | | | 3.09 | 29 | -75 | 22 | Transverse occipital sulcus |
| | | | 3.09 | 32 | 22 | 3 | Anterior insula |
| | | | 3.09 | 27 | -86 | -3 | Middle occipital gyrus |
| | | | 3.09 | 26 | -32 | -13 | Parahippocampal gyrus |
| | | | 3.09 | 26 | -1 | -16 | Parahippocampal gyrus |
| | | | 3.09 | 22 | -76 | 31 | Superior parietal lobule |
| *Word class effects: actions > objects in fMRI1 (S1_VU + S1_VT > S1_NU + S1_NT)* | | | | | | | |
| LH | 2.80 | 5871 | 3.09 | -39 | -74 | 11 | Middle occipital gyrus |
| | | | 3.09 | -42 | -54 | 12 | Superior temporal sulcus |
| | | | 3.09 | -43 | -74 | -2 | Inferior occipital gyrus |
| | | | 3.09 | -46 | -55 | 8 | Superior temporal sulcus |

| | | | | | | | |
|---|---|---|---|---|---|---|---|
| | | | 3.09 | -44 | -65 | 12 | Middle temporal gyrus |
| | | | 3.09 | -48 | -62 | 2 | Middle temporal gyrus |
| | | | 3.09 | -51 | -46 | 8 | Superior temporal sulcus |
| | | | 3.09 | -55 | -55 | 0 | Middle temporal gyrus |
| LH | 2.18 | 1282 | 2.29 | -31 | -47 | 47 | Superior parietal lobule |
| LH | 2.15 | 806 | 2.29 | -39 | -46 | -14 | Fusiform gyrus |
| LH | 2.04 | 160 | 2.12 | -21 | -83 | 30 | Superior parietal lobule |
| LH | 2.00 | 110 | 2.01 | -48 | -27 | 34 | Intraparietal sulcus |
| LH | 1.99 | 72 | 2.00 | -25 | -70 | 22 | Superior parietal lobule |
| LH | 1.99 | 15 | 2.00 | -29 | -53 | 36 | Superior parietal lobule |
| LH | 1.99 | 33 | 2.00 | -55 | -25 | 30 | Intraparietal sulcus |
| RH | 2.66 | 8200 | 3.09 | 50 | -50 | 4 | Middle temporal gyrus |
| | | | 3.09 | 44 | -62 | -1 | Middle temporal gyrus |
| | | | 3.09 | 48 | -51 | 10 | Middle temporal gyrus |
| | | | 3.09 | 42 | -66 | 12 | Middle occipital gyrus |
| | | | 3.09 | 39 | -75 | 10 | Middle occipital gyrus |
| | | | 3.09 | 42 | -58 | 8 | Middle temporal gyrus |
| | | | 3.09 | 38 | -55 | -12 | Fusiform gyrus |
| | | | 3.09 | 42 | -42 | -13 | Fusiform gyrus |
| | | | 2.65 | 46 | -43 | 14 | Superior temporal sulcus |
| RH | 1.97 | 54 | 1.98 | 26 | -90 | -7 | Inferior occipital gyrus |
| *Word class effects: objects > actions in fMRI 1 (S1_NU + S1_NT > S1_VU + S1_VT)* | | | | | | | |
| LH | 2.17 | 498 | 2.46 | -7 | -82 | -9 | Fusiform gyrus |
| *Training effects: trained > untrained nouns in fMRI2 (S2_NT > S2_NU)* | | | | | | | |
| LH | -2.02 | 241 | -2.10 | -29 | 15 | 5 | Anterior insula |
| LH | -2.02 | 821 | -2.05 | -45 | 11 | 17 | Inferior frontal gyrus (pars opercularis) |
| *Training effects: trained > untrained verbs in fMRI2 (S2_VT > S2_VU)* | | | | | | | |
| LH | -2.46 | 2588 | -2.88 | -42 | 9 | 23 | Inferior frontal gyrus (pars opercularis) |
| LH | -2.42 | 2419 | -2.88 | -35 | 23 | 5 | Anterior insula |
| *Session effects: untrained nouns in fMRI2 > untrained nouns in fMRI1 (S2_NU > S1_NU)* | | | | | | | |
| LH | -2.09 | 962 | -2.37 | -16 | -51 | -4 | Fusiform gyrus |
| LH | -2.05 | 203 | -2.14 | -17 | -68 | -8 | Fusiform gyrus |
| LH | -2.04 | 98 | -2.10 | -22 | -14 | 50 | Superior frontal sulcus |
| LH | -1.99 | 51 | -2.01 | -20 | -62 | 3 | Calcarine sulcus |
| LH | -2.01 | 142 | -2.01 | -17 | -70 | 48 | Superior parietal lobule |
| LH | -1.98 | 29 | -2.00 | -4 | -62 | 2 | Fusiform gyrus |
| RH | -2.16 | 3442 | -2.26 | 12 | -55 | 4 | Calcarine sulcus |
| RH | -2.07 | 211 | -2.14 | 19 | -62 | 48 | Superior parietal lobule |
| *Session effects: untrained verbs in fMRI2 > untrained verbs in fMRI1 (S2_VU > S1_VU)* | | | | | | | |
| LH | -2.04 | 68 | -2.07 | -15 | -72 | -8 | Fusiform gyrus |
| RH | -2.08 | 889 | -2.14 | 14 | -65 | 49 | Superior parietal lobule |

| | | | | | | | |
|---|---|---|---|---|---|---|---|
| RH | -2.04 | 2380 | -2.12 | 12 | -59 | 8 | Calcarine sulcus |
| RH | -1.98 | 108 | -2.01 | 15 | -71 | 40 | Superior parietal lobule |
| *Training effects > Session effects for nouns ((S2_NT > S1_NT) > (S2_NU > S1_NU))** | | | | | | | |
| LH | -1.68 | 86 | -1.71 | -46 | 13 | 11 | Inferior frontal gyrus (frontal operculum) |
| LH | -1.67 | 87 | -1.70 | -43 | 15 | 25 | Inferior frontal gyrus (pars triangularis) |
| LH | -1.66 | 7 | -1.68 | -49 | 12 | 20 | Inferior frontal gyrus (pars triangularis) |
| *Training effects > Session effects for verbs ((S2_VT > S1_VT) > (S2_VU > S1_VU))** | | | | | | | |
| LH | -1.89 | 1222 | -2.26 | -40 | 8 | 26 | Inferior frontal gyrus (pars opercularis) |
| LH | -1.85 | 772 | -2.05 | -29 | 16 | 3 | Anterior insula |
| LH | -1.79 | 444 | -1.98 | -43 | 33 | 8 | Inferior frontal gyrus (pars orbitalis) |

**Supplementary Table 2. Clusters identified in the whole-brain multivariate (searchlight) analyses that showed decoding accuracy significantly greater than chance.** To obtain cluster statistics, we projected the surface maps back into the volume. The table shows average *z*-values and extent (in $mm^3$) of active clusters, as well as the Talairach coordinates of each cluster's center-of-gravity (COG), its corresponding *z*-value and anatomical label. For clusters larger than 5000 $mm^3$ local maxima were identified using the NeuroElf function *clustervol.m*. Clusters with volume smaller than 10 $mm^3$ are not reported. Maps were thresholded at $z > 1.96$ (two-tailed). LH = left hemisphere, RH = right hemisphere.

| cluster | | | local maxima | | | | |
|---|---|---|---|---|---|---|---|
| **side** | **mean *Z*** | **$mm^3$** | ***Z*** | $x_{COG}$ | $y_{COG}$ | $z_{COG}$ | **anatomical region** |
| *Objects vs. actions in fMRI1* | | | | | | | |
| LH | 2.77 | 53629 | 3.09 | -5 | -76 | 10 | Cuneus |
| | | | 3.09 | -7 | -79 | 2 | Calcarine sulcus |
| | | | 3.09 | -7 | -64 | -2 | Fusiform gyrus |
| | | | 3.09 | -10 | -75 | -10 | Fusiform gyrus |
| | | | 3.09 | -11 | -86 | 17 | Cuneus |
| | | | 3.09 | -10 | -69 | 28 | Parieto-occipital sulcus |
| | | | 3.09 | -17 | -52 | -5 | Fusiform gyrus |
| | | | 3.09 | -17 | -82 | -11 | Fusiform gyrus |
| | | | 3.09 | -14 | -75 | 38 | Parieto-occipital sulcus |
| | | | 3.09 | -21 | -77 | 27 | Superior parietal lobule |
| | | | 3.09 | -26 | -88 | 5 | Middle occipital gyrus |
| | | | 3.09 | -23 | -59 | -10 | Fusiform gyrus |
| | | | 3.09 | -26 | -71 | 30 | Superior parietal lobule |
| | | | 3.09 | -26 | -63 | 44 | Superior parietal lobule |
| | | | 3.09 | -29 | -79 | 19 | Superior occipital gyrus |
| | | | 3.09 | -33 | -48 | -14 | Fusiform gyrus |
| | | | 3.09 | -29 | -50 | 48 | Superior parietal lobule |
| | | | 3.09 | -34 | -74 | -12 | Fusiform gyrus |
| | | | 3.09 | -38 | -39 | -14 | Fusiform gyrus |
| | | | 3.09 | -38 | -70 | 26 | Angular gyrus |
| | | | 3.09 | -40 | -71 | 10 | Middle occipital gyrus |
| | | | 3.09 | -43 | -55 | 17 | Superior temporal gyrus |
| | | | 3.09 | -40 | -36 | 44 | Superior parietal lobule |
| | | | 3.09 | -41 | -68 | -8 | Inferior occipital gyrus |
| | | | 3.09 | -50 | -53 | -6 | Middle temporal gyrus |
| | | | 3.09 | -47 | -65 | 4 | Middle temporal gyrus |
| | | | 3.09 | -51 | -43 | 4 | Middle temporal gyrus |
| | | | 3.09 | -52 | -32 | 27 | Supramarginal gyrus |
| | | | 3.09 | -52 | -46 | 16 | Supramarginal gyrus |
| | | | 3.09 | -56 | -33 | 9 | Superior temporal gyrus |
| | | | 2.65 | -8 | -50 | 38 | Precuneus |

| | | | | | | | |
|---|---|---|---|---|---|---|---|
| LH | 2.25 | 6310 | 2.51 | -42 | 17 | 22 | Inferior frontal gyrus (pars triangularis) |
| | | | 2.51 | -46 | 2 | 26 | Precentral sulcus |
| | | | 2.51 | -49 | -1 | 12 | Inferior frontal gyrus (frontal operculum) |
| | | | 2.46 | -41 | 11 | 23 | Inferior frontal gyrus (pars triangularis) |
| | | | 2.46 | -37 | 2 | 46 | Middle frontal gyrus |
| | | | 2.41 | -47 | -7 | 38 | Precentral gyrus |
| LH | 2.03 | 212 | 2.12 | -38 | 16 | 36 | Middle frontal gyrus |
| RH | 2.82 | 44008 | 3.09 | 52 | -37 | 7 | Superior temporal sulcus |
| | | | 3.09 | 48 | -56 | 2 | Middle temporal gyrus |
| | | | 3.09 | 45 | -43 | -14 | Inferior temporal gyrus |
| | | | 3.09 | 44 | -57 | 20 | Middle temporal gyrus |
| | | | 3.09 | 36 | -52 | -8 | Fusiform gyrus |
| | | | 3.09 | 38 | -75 | 10 | Middle occipital gyrus |
| | | | 3.09 | 33 | -60 | -10 | Fusiform gyrus |
| | | | 3.09 | 35 | -70 | -11 | Inferior occipital gyrus |
| | | | 3.09 | 28 | -63 | 38 | Superior parietal lobule |
| | | | 3.09 | 33 | -80 | 2 | Middle occipital gyrus |
| | | | 3.09 | 26 | -75 | 24 | Superior parietal lobule |
| | | | 3.09 | 28 | -38 | -14 | Fusiform gyrus |
| | | | 3.09 | 26 | -78 | -11 | Inferior occipital gyrus |
| | | | 3.09 | 20 | -89 | 1 | Middle occipital gyrus |
| | | | 3.09 | 12 | -83 | 4 | Calcarine sulcus |
| | | | 3.09 | 15 | -74 | 31 | Parieto-occipital sulcus |
| | | | 3.09 | 10 | -67 | -4 | Fusiform gyrus |
| | | | 3.09 | 8 | -82 | 13 | Cuneus |
| | | | 2.88 | 32 | -41 | 46 | Superior parietal lobule |
| | | | 2.75 | 7 | -58 | 29 | Precuneus |
| RH | 2.18 | 387 | 2.41 | 48 | 23 | 22 | Inferior frontal gyrus (pars triangularis) |
| RH | 2.04 | 15 | 2.12 | 53 | 27 | 15 | Inferior frontal gyrus (pars triangularis) |
| RH | 2.02 | 31 | 2.12 | 46 | 8 | 25 | Inferior frontal gyrus (pars opercularis) |
| *Trained vs. untrained items in fMRI2* | | | | | | | |
| LH | 2.38 | 489 | 3.09 | -39 | -63 | 20 | Superior temporal sulcus |
| LH | 2.31 | 98 | 2.58 | -4 | -65 | 28 | Precuneus |
| LH | 2.31 | 84 | 2.58 | -28 | 24 | 5 | Anterior insula |
| LH | 2.26 | 139 | 2.51 | -43 | -57 | 39 | Angular gyrus |
| LH | 2.16 | 67 | 2.29 | -28 | -75 | 26 | Angular gyrus |
| LH | 2.18 | 22 | 2.29 | -36 | -61 | 40 | Angular gyrus |
| LH | 2.12 | 29 | 2.29 | -41 | -64 | 37 | Angular gyrus |
| LH | 2.14 | 30 | 2.26 | -36 | -71 | 36 | Angular gyrus |
| LH | 2.11 | 11 | 2.23 | -7 | -68 | 23 | Precuneus |
| RH | 2.15 | 44 | 2.33 | 3 | -77 | -6 | Fusiform gyrus |
